# Delayed effects of climate on vital rates lead to demographic divergence in Amazonian forest fragments

**DOI:** 10.1101/2021.06.28.450186

**Authors:** Eric R. Scott, María Uriarte, Emilio M. Bruna

**Author notes:** Correspondence: Eric R. Scott < >.

## Abstract

Deforestation often results in landscapes where remaining forest habitat is highly fragmented, with remnants of different sizes embedded in an often highly contrasting matrix. Local extinction of species from individual fragments is common, but the demographic mechanisms underlying these extinctions are poorly understood. It is often hypothesized that altered environmental conditions in fragments drive declines in reproduction, recruitment, or survivorship. The Amazon basin, in addition to experiencing continuing fragmentation, is experiencing climate change related increases in the frequency and intensity of droughts and unusually wet periods. Whether plant populations in tropical forest fragments are particularly susceptible to extremes in precipitation remains unclear. Most studies of plants in fragments are relatively short (1–6 years), focus on a single life-history stage, and often do not compare to populations in continuous forest. Even fewer studies consider delayed effects of climate on demographic vital rates despite the importance of delayed effects in studies that consider them. Using a decade of demographic and climate data from an experimentally fragmented landscape in the Central Amazon, we assess the effects of climate on populations of an understory herb (*Heliconia acuminata*, Heliconiaceae). We used distributed lag non-linear models to understand the delayed effects of climate (measured as standardized precipitation evapotranspiration index, SPEI) on survival, growth, and flowering. We detected delayed effects of climate up to 36 months. Extremes in SPEI in the previous year reduced survival, drought in the wet season 8–11 months prior to the February census increased growth, and drought two dry seasons prior increased flowering probability. Effects of extremes in precipitation on survival and growth were more pronounced in forest fragments compared to continuous forest. The complex delayed effects of climate and habitat fragmentation in our study point to the importance of long-term demography experiments in understanding the effects of anthropogenic change on plant populations.

## Introduction

The expansion of agriculture and other human activities is a primary driver of deforestation in the tropics (Alroy, 2017; Haddad et al., 2015). It also results in landscapes where the remaining forest can be highly fragmented, with patches of different sizes embedded in a matrix of often contrasting habitat (Bianchi & Haig, 2013; Taubert et al., 2018). This fragmentation is associated with myriad ecological changes, including the local and regional extinction of plant species (da Silva & Tabarelli, 2000; Laurance et al., 2006). It is often hypothesized that the dramatically altered environmental conditions in tropical forest fragments (Arroyo-Rodríguez et al., 2017; Didham & Lawton, 1999; Ewers & Banks-Leite, 2013) drive declines in plant reproduction, recruitment, or survivorship (Bruna, 1999; Laurance et al., 1998; Zartman et al., 2015), although the demographic mechanisms responsible for these extinctions are poorly understood (Bruna et al., 2009). Despite the prevalence of this hypothesis (Betts et al., 2019; Didham & Lawton, 1999; Laurance et al., 2001), efforts to link population-level demographic responses with altered environmental conditions in fragments remains scarce.

Studies in temperate systems have shown that the demography of species can also be altered by climate change (Doak & Morris, 2010; Selwood et al., 2015; Sletvold, 2005; Williams et al., 2015). While the demographic consequences of climate change for tropical species are expected to be similarly severe (Brodie et al., 2012; Scheffers et al., 2017), surprisingly little is known about the responses of these species to climatic variability (Paniw et al., 2021). Climate models predict increases in the areas of the Amazon effected by severe drought and unusual wetness by the year 2100 (Duffy et al., 2015). Tropical plants may be particularly sensitive to climate change—they typically have narrow ranges of climatic tolerance (Feeley et al., 2012), and recent results suggest increases in the frequency and severity of extreme precipitation events (both drought and extreme wet) reduce survival and reproduction (Esteban et al., 2021; Gaoue et al., 2019). This sensitivity to climatic fluctuations, coupled with evidence that plant growth and survivorship are lower in fragments (Bruna et al., 2002; Laurance et al., 1998; Zartman et al., 2015), has led to speculation that plants in forest fragments will be especially susceptible to climate change (Laurance et al., 2001; Opdam & Wascher, 2004; Selwood et al., 2015).

The simultaneous pressures of climate change and habitat fragmentation may also result in worse than additive impacts on demography (Holyoak & Heath, 2016; Oliver et al., 2015). For example, if fragments have a reduced capacity to buffer changes in microclimate (Didham & Lawton, 1999; Ewers & Banks-Leite, 2013) or if fragment connectivity is climate dependent (Honnay et al., 2002), populations experiencing habitat fragmentation will fare worse under climate change.

Whether the demography of plant populations in tropical forest fragments is more susceptible to climatic extremes remains unclear for three primary reasons. First, most studies of plants in fragments have focused on a single life-history stage or process (Bruna et al., 2009; Ehrlen et al., 2016), making it challenging to draw broader demographic conclusions. Second, there is a growing literature on how tropical plants respond to droughts (Esquivel-Muelbert et al., 2019; González-M et al., 2020; Uriarte et al., 2016), but few studies have compared the responses of plants in continuous forest with those of plants in forest fragments (Laurance et al., 2001). Finally, the multi-year data needed to test population-level hypotheses about climate change and fragmentation are scant, especially for tropical systems (Crone et al., 2011; Salguero-Gómez et al., 2015). These data are critical not simply because they allow for capturing variation in climatic conditions and the resulting demographic responses (Morris & Doak, 2002; Teller et al., 2016). They are also essential because while some demographic effects of fragmentation or drought can be detected immediately, others may take years to manifest (*e.g.,* Gagnon et al., 2011). Indeed, lagged responses of demographic vital rates to climate may be the rule rather than the exception (Anderegg et al., 2015; Evers et al., 2021; Kannenberg et al., 2020; Schwalm et al., 2017).

Herbaceous plants represent up to 25% of plant diversity in tropical forests (Gentry & Dodson, 1987), are critical food and habitat for myriad species (Snow, 1981), and are economically and culturally vital (Nakazono et al., 2004; Ticktin, 2003). Nevertheless, the impacts of global change phenomena on their demography remain conspicuously understudied (Bruna et al., 2009). We used a decade of demographic and climatic data from an experimentally fragmented landscape in the Central Amazon to assess the effects of climate on populations of a tropical understory herb (*Heliconia acuminata*, Heliconiaceae). This time series, which included the severe droughts of 1997 (McPhaden, 1999) and 2005 (Marengo et al., 2008; Zeng et al., 2008), allowed us to address the following questions: (1) Does drought increase or decrease the growth, survival, and fertility of plant populations in continuous forest? (2) Are there delayed effects of drought on demographic vital rates, and if so what lag times are most critical? (3) Are the effects of drought on the vital rates of populations in fragments similar in direction and magnitude to those in continuous forest?

## Methods

### Study site

The Biological Dynamics of Forest Fragments Project (BDFFP) is located ∼70 km north of Manaus, Brazil (2°30’ S, 60°W). In addition to large areas of continuous forest, the BDFFP has forest fragment reserves isolated from 1980–1984 by felling the trees surrounding the area chosen for isolation and, in most cases, burning the downed trees once they dried (Bierregaard et al., 1992). In subsequent decades the vegetation regenerating around fragments has been periodically cleared to ensure fragment isolation (Bierregaard et al., 2001).

The BDFFP reserves are located in nonflooded (i.e., *terra firme*) tropical lowland forest with a 30–37m tall canopy (Rankin-de-Mérona et al., 1992) and an understory dominated by stemless palms (Scariot, 1999). The soils in the reserves are nutrient-poor xanthic ferrosols; their water retention capacity is poor despite having a high clay content. Mean annual temperature in the region is 26° C (range=19–39° C), and annual rainfall ranges from 1900–2300 mm. There is a pronounced dry season from June to October (Figure S1).

### Focal species

*Heliconia acuminata* (LC Rich.) (Heliconiaceae) is a perennial monocot distributed throughout Central Amazonia (Kress, 1990) and is the most abundant understory herb at the BDFFP (Ribeiro et al., 2010). While many *Heliconia* species grow in large patches in treefall gaps and other disturbed areas, others, such as *H. acuminata,* are found at lower densities in the darker and cooler forest understory (Rundel et al., 2020). These species produce fewer inflorescences and are pollinated by traplining rather than territorial hummingbirds (Bruna et al., 2004; Stouffer & Bierregaard, 1996). In our sites *H. acuminata* is pollinated by *Phaeothornis superciliosus* and *P. bourcieri* (Bruna et al., 2004). Plants begin flowering at the start of the rainy season; reproductive plants have *x̄* = 1.1 flowering shoots (range = 1–7), each of which has an inflorescence with 20–25 flowers (Bruna & Kress, 2002). Fruits mature April-May, have 1–3 seeds per fruit (*x̄* = 2), and are eaten by a thrush and several species of manakin (Uriarte et al., 2011). Dispersed seeds germinate approximately 6 months after dispersal at the onset of the subsequent rainy season, with rates of germination and seedling establishment higher in continuous forest than forest fragments (Bruna, 1999; Bruna & Kress, 2002).

### Demographic data collection

This study uses data collected in four 1-ha fragment reserves and six continuous forest sites. In 1997–1998 we established a 5000 m^2^ plots (50 × 100m) in each of these sites in which we marked and measured all *Heliconia acuminata*; plots in 1-ha fragments were on one randomly selected half of the fragment, while plots in continuous forest were located 500–4000 m from the borders of secondary and mature forest. The distance between plots ranged from 500 m–41 km. Our dataset comprised 4,886 plants in continuous forest and 1,375 plants in forest fragments. Plots in CF had on average more than twice as many plants than plots in 1-ha fragments (CF median = 788, range = [201, 1549]; 1-ha median = 339, range = [297, 400]).

Each plot was subdivided into 50 quadrats (10 × 10m) to simplify annual surveys, during which we recorded the number of vegetative shoots each plant had, the height of each plant (i.e. distance from the ground to the tallest leaf tip), and whether each plant was flowering (height and shoot number are correlated with leaf area, the probability of flowering, and rates of survivorship (Bruna, 2002; Bruna & Kress, 2002). In this study, we used the product of shoot number and plant height as our measure of plant size. Preliminary analysis showed that the product of shoot number and height was a better predictor of total leaf area (which in turn is assumed to be a strong predictor of aboveground biomass) than either shoot number or height alone (Table S2). After the initial census year, between 80% and 97% of marked plants were found at each survey. Of plants that had missing values for some years, but were found again in a subsequent year, 95% had 2 or fewer years of missing values. Therefore, plants that were not found for three consecutive surveys, and no subsequent survey, were considered to have died in the transition year after their last observation. We also surveyed plots regularly during the rainy season to identify any that flowered after the survey. For additional details on the location of plots, survey methods, and *H. acuminata* population structure see Bruna & Kress (2002).

### Climate data

Data on precipitation and potential evapotranspiration in our sites were obtained from a published gridded dataset (0.25° × 0.25° resolution) built using data from 3,625 ground-based weather stations across Brazil (Xavier et al., 2016). We used these data to calculate the standardized precipitation evapotranspiration index (SPEI) (Vicente-Serrano et al., 2010). SPEI is a proxy for meteorological drought that integrates precipitation and evapotranspiration anomalies over a specified time scale. Positive SPEI values for a given month indicate conditions wetter than the historical average for that month, while negative values of SPEI indicate droughts with intensity categorized as mild (0 to -1), moderate (-1 to -1.5), severe (-1.5 to -2), or extreme (< -2) (McKee et al., 1993). SPEI can be calculated to represent different temporal scales of drought; we used 3-month SPEI because—given its shallow roots and rhizome—*H. acuminata* relies primarily on soil moisture rather than deeper water sources that can persist for longer timescales (McKee et al., 1993; Vicente-Serrano et al., 2010). Note that 3-month SPEI is still monthly data—each month’s SPEI value simply takes into account precipitation and evapotranspiration of the previous three months. SPEI calculations were made using the `SPEI` package in R (Beguería & Vicente-Serrano, 2017). The timing of drought events based on these SPEI calculations is consistent with that resulting from SPEI calculated with other data sources, though the magnitude of drought sometimes differed (Figure S2; Table S1).

### Statistical Modeling of Vital Rates

To assess the effects of drought history on plant vital rates we used Distributed Lag Non-linear Models (DLNMs, Gasparrini et al., 2017). DLNMs capture how potentially delayed effects of predictor variables (e.g. SPEI) affect an outcome (e.g. growth) well beyond the event period. DLNMs avoid the problems of including all possible climate windows in a single model—namely collinearity of predictors and the arbitrary choice of window length. They do so by fitting a bi-dimensional predictor-lag-response association spline, referred to as a crossbasis function. This models a non-linear relationship between predictor and response (e.g. between SPEI and vital rates) and allows the shape of that relationship to vary smoothly over lag time. The shape of the function can be penalized so that it is only as complex as the data supports, thus reducing effective degrees of freedom. We used a crossbasis function of SPEI and lag time with possible lags from 0–36 months. We chose 36 months as a maximum lag because prior transplant experiments with *H. acuminata* showed they typically recover from transplant shock in less than 36 months (Bruna et al., 2002) so this is a reasonable upper bound for lagged effects of drought. The general form of the vital rate (*y*) models was as follows:

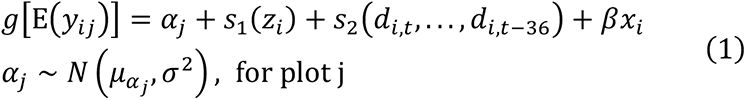

where *y_ij_* is the *i*th plant observation in the *j*th plot *s*_1_(*z_i_*) is a smooth function of plant size (natural log of height × shoot number), fit using a penalized cubic regression spline, *s*_2_(⋅) is the crossbasis function in which *d_i,t_* is the SPEI value during the census month of an observation (February) and *d_i,t-1_* is the SPEI *l* months prior (see Gasparrini et al. 2017 for details). To determine if plot characteristics influenced average vital rates we included a random effect of plot ID on the intercept; this was represented by *a_j_* in eq. 1. We modeled a potential cost of reproduction by including flowering in the previous year as covariate, *x_i_*, in eq. 1. Individual-level random effects were not included due to computational limitations in fitting random intercepts for all of the nearly 5,000 plants in continuous forest plots. Fitting models with a data subset with and without individual-level random effects showed that excluding individual-level random effects reduced model R^2^, but did not affect the shape or statistical significance of the smooths.

The crossbasis function, s_2_ (⋅) can also be written:

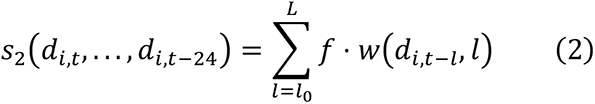

where the crossbasis function, *f* ⋅ *w*(*d*, *l*), is composed of two marginal basis functions: the standard predictor-response function *f*(*d*), and the additional lag-response function *w*(*l*). These marginal functions are combined as a tensor product smooth such that the shape of one marginal function varies smoothly along the other dimension (see chapter 5 of Wood (2017) and Gasparrini et al. (2017) for more detail). Penalized cubic regression splines were used for both marginal bases of the crossbasis function. We increased *k*, the maximum basis dimension, as necessary to be sure it was adequate (Appendix A). Because of penalization, the maximum basis dimension is generally not important as long it is large enough to allow the smooth to represent the ‘true’ relationship (Wood, 2017). Estimated degrees of freedom (edf) represent the ‘true’ complexity of the smooth after penalization with edf = 1 being equivalent to a straight line and larger numbers representing more complex curves.

The crossbasis function was fit to the data in the context of a generalized additive model (GAM) with restricted maximum likelihood using the `mgcv` R package (Wood, 2017). The crossbasis smooth was coded as:

~~~
s(spei_history, L, bs = "cb", xt = list(bs = "cr"))
~~~

Where `spei_history` is a *n* × *m* matrix with each row (*n*) containing the climate history of a single plant in a year and *m* = 36 columns representing the past weather conditions starting at the month of the census and going backwards in time 36 months. `L` is an *n* × *m* matrix with columns describing the lag structure, i.e. the integers 0 through 36. The spline basis, `“cb”` is provided by the `dlnm` package (Gasparrini, 2011), and we specified the marginal bases as cubic regression splines with the `xt` argument. For the full code used to fit the models, see the Data Availability Statement.

We determined the effects of SPEI on plant growth using plant size in year t+1 as a response variable. This was modeled with a scaled t family error distribution because residuals were leptokurtic when a Gaussian family was used. Because number of inflorescences was highly zero-inflated, we converted this to a binary response to model reproduction (i.e., 1 for ≥1 inflorescence, 0 for no inflorescences). We modeled both reproduction and survival (i.e., from year t to year t+1) using a binomial family error distribution with a logit link function.

In the process of fitting the models, the penalty on the crossbasis smooth (and other smoothed terms) is optimized such that more linear shapes are favored unless the data supports non-linearity (Wood, 2017). We applied an additional penalty to shrink linear portions toward zero with the `select=TRUE` option to the `gam()` function, and inferred statistical significance of model terms with p-values from the `summary.gam()` function as recommended in Marra & Wood (2011).

The `dlnm` package does not currently allow the modeling of interaction terms, which means we could not asses the interaction of habitat type and lagged effects. We therefore fit separate models for plants in fragments and in continuous forest to allow the shape of the crossbasis function to differ between habitats. Significant main effects of habitat type were assessed by looking for overlap in the 84% confidence intervals of model intercepts; the 84% CIs of two samples drawn from the same population overlap about 95% of the time (Payton et al., 2003).

To visualize results, we plotted partial effects plots for the one-dimensional smooth function of log size in the previous year and the two-dimensional smooth function of lagged SPEI (the crossbasis function) for each model. To make these plots more interpretable, we added the model intercept and (when applicable) back-transformed the y-axis to be on the scale of the response (e.g. a probability for survival and flowering).

All analyses were conducted in R version 4.0.2 (2020-06-22) (R Core Team, 2020) using the `targets` package for workflow management (Landau, 2021). Figures were created with the aid of the `gratia`, `ggplot2`, and `patchwork` packages (Pedersen, 2020; Simpson, 2021; Wickham, 2016).

## Results

The meteorological droughts in our focal region indicated by SPEI are generally consistent with those reported in the literature (Table S1). For example, the drought associated with the 1997 El Niño Southern Oscillation (ENSO) event was one of the most severe on record for the Amazon (Williamson et al., 2000); correspondingly, 1997 has the lowest SPEI values in our timeseries (Figure 1d). The 2005 dry season (June–October) was also reported as an exceptionally dry year, although this drought mostly affected the southwestern Amazon (Marengo et al., 2008; Zeng et al., 2008). Our SPEI data show the 2005 dry season to be a moderate drought (-1 > SPEI > -1.5).

**Figure 1:**
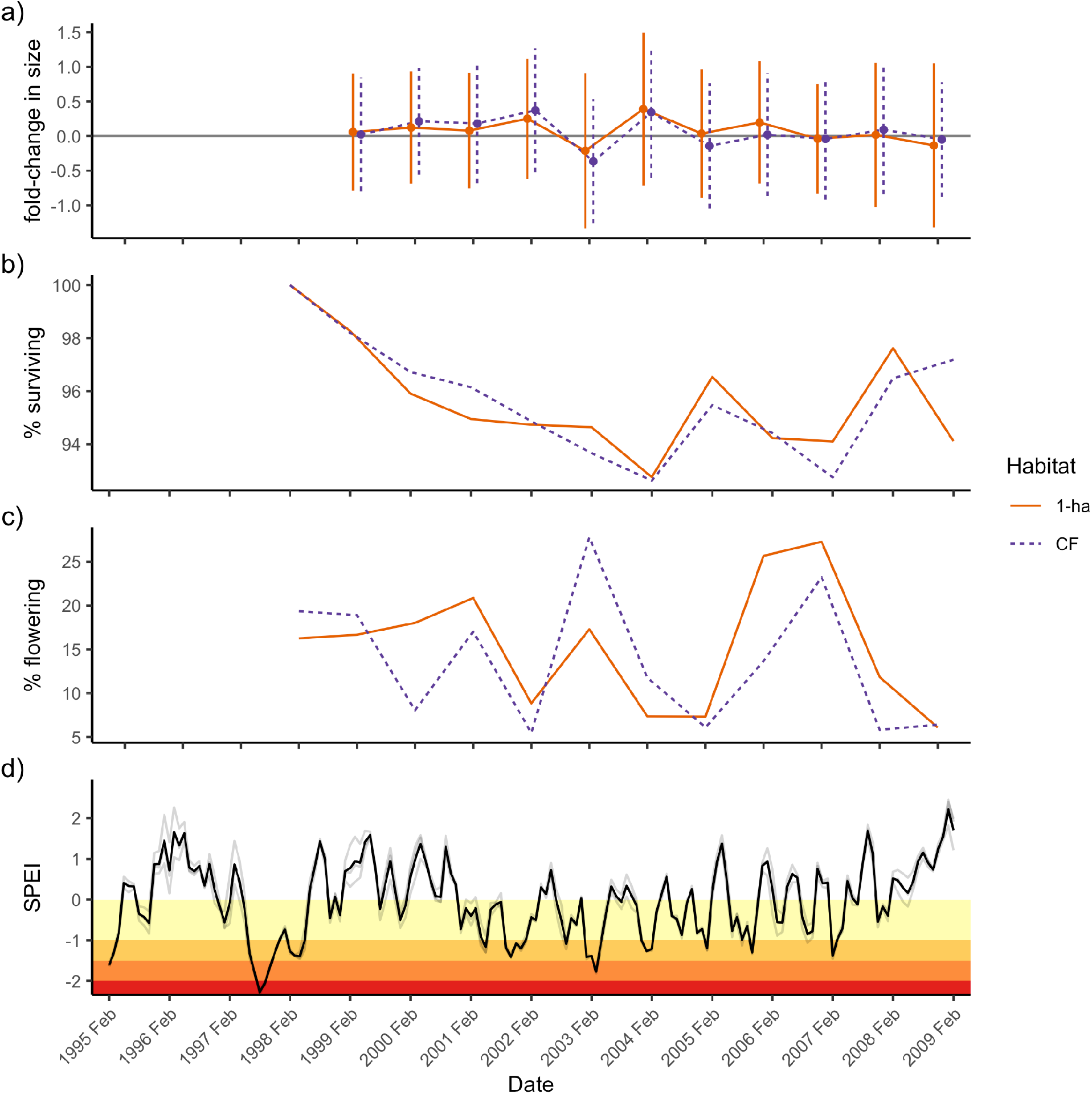
(a-c) Time series of *H. acuminata* vital rates in 1-ha fragments (solid orange lines) and continuous forest (dashed blue lines) and (d) drought occurrence in the study region. (a) Mean fold-change in plant plant size (log2(size_t+1_ / size_t_)) varies by year and habitat. On average, plants grew in most years with the notable exception in 2003, in which on average plants regressed in size in both habitats (i.e., fold-change < 0). Error bars represent the standard deviation. (b) The proportion of plants surviving from one transition year to the next varied from 0.98 (CF in 1998-1999) to 0.93 (CF in 2003-2004). (c) The proportion of *H. acuminata* above the size threshold for reproduction that flowered each year is on average low but variable. The size threshold is determined by the upper 90th percentile size of flowering plants across all years. (d) Monthly 3-month SPEI for our study region. Gray lines represent values from different grid cells encompassing BDFFP; the dark line represents the site mean. Colored stripes represent drought intensity: yellow = mild, orange = moderate, dark orange = severe, red = extreme.

### Survival, size, and flowering in continuous forest vs. fragments

#### Survival

Across all plots, the proportion of plants surviving was lowest in the 2003–2004 transition year (93% survival in both continuous forest and forest fragments). This coincided with droughts in both the 2003 and 2004 rainy seasons (Figure 1b,d) and was preceded by a drop in average plant size in the 2002–2003 transition year (i.e. negative growth in Figure 1a). There was no significant difference in survival between continuous forest and 1 ha fragments (Table 1). Although differences in survival probability are small and not statistically significant, these slight differences can be seen to compound over time in the survivorship of plants labeled in the first year of the surveys (Figure 2). Survival in both habitats was significantly affected by size in the previous year (Table 2) with larger plants having much greater survival probabilities (Figure 3b). The survival probability of large plants approached 1 in both habitat types, but the smallest plants had higher survival in 1 ha fragments (although 95% confidence intervals overlap at all sizes).

**Table 1:** Model intercepts and their 84% confidence intervals. Non-overlapping confidence intervals are bolded and can be interpreted as a significant difference in the vital rate between continuous forest and 1 ha fragments.

| vital rate | habitat | intercept |
| --- | --- | --- |
| survival | CF | 0.97 [0.964, 0.975] |
|  | 1 ha | 0.962 [0.957, 0.966] |
| size | CF | **4.36 [4.313, 4.408]** |
|  | 1 ha | **4.18 [4.062, 4.297]** |
| flowering | CF | **0.006 [0.005, 0.009]** |
|  | 1 ha | **0.002 [0.001, 0.004]** |

**Figure 2:**
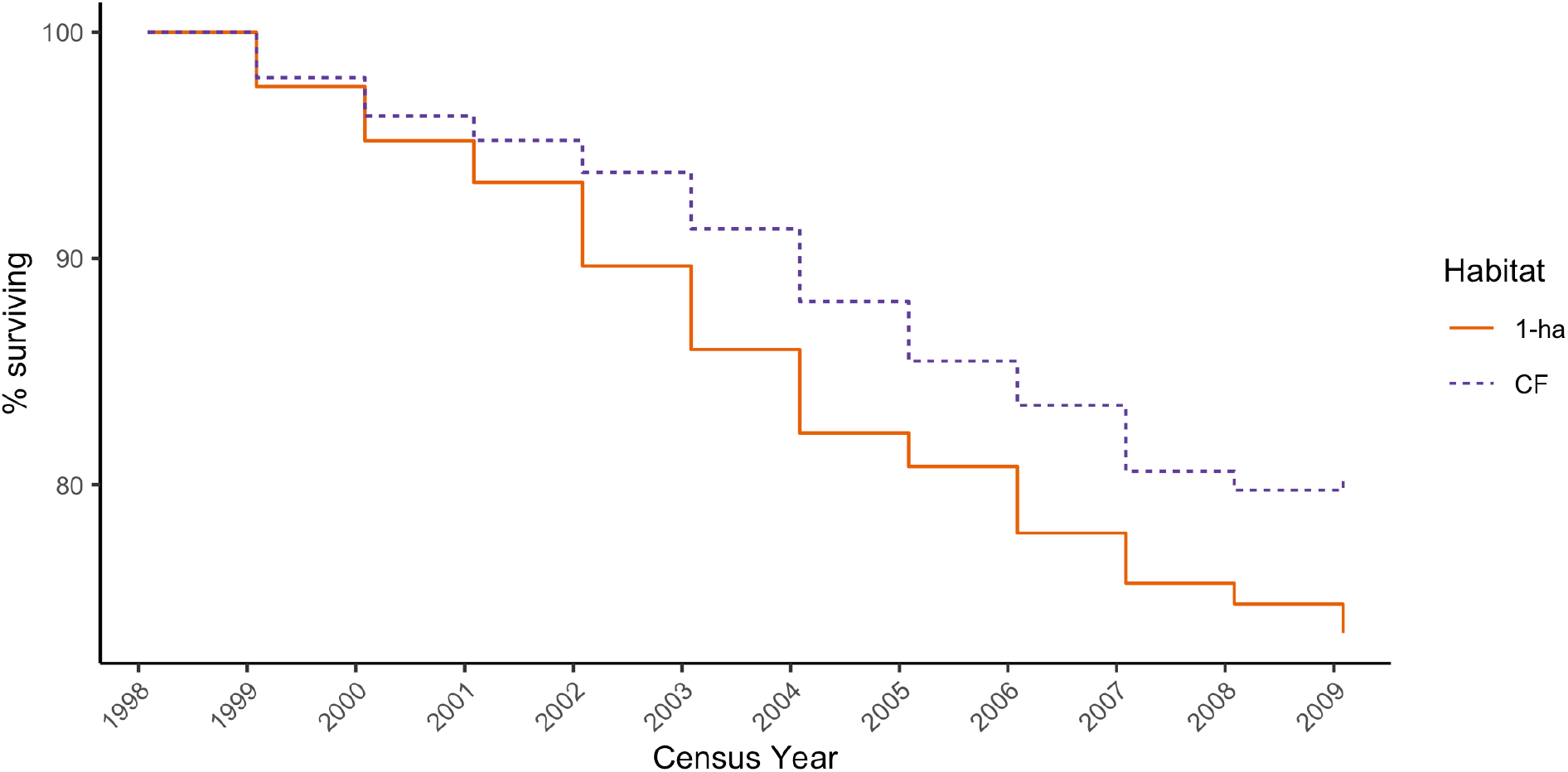
Survivorship curve for plants marked in the 1998 survey year; these plants comprise 49% of those in the complete demographic dataset. The percentage of these plants that were still alive ten years later was 79.7% (1629/2055) in continuous forest vs. 72.4% (393/543) in 1-ha fragments.

**Table 2:**
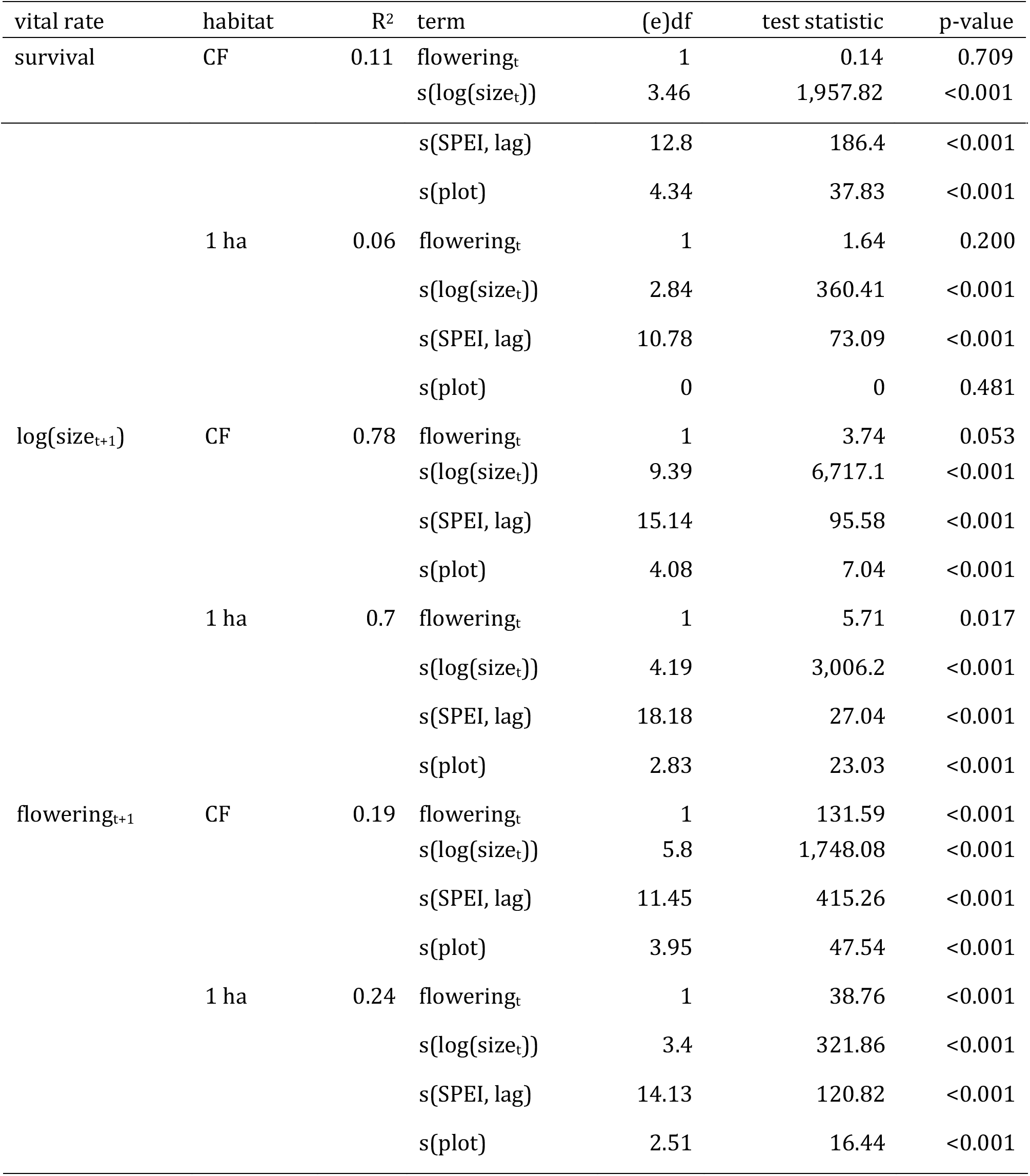
Marginal hypothesis tests for models of survival probability, size, and flowering probability of *H. acuminata* in continuous forest and 1 ha fragment plots. The adjusted R^2^ is reported as a measure of model fit. The model terms included the parametric fixed effect factor of whether plants flowered the previous year (“flowering_t_”), the smoothed fixed effect of plant size (“s(log(size_t_))”), the crossbasis smooth of lagged SPEI (“s(SPEI, lag)”), and a random effect of plot ID, (“s(plot)”). Degrees of freedom are reported for the parametric term, and estimated degrees of freedom (edf) are reproted for smooths. Larger values for edf indicate more complex smooths and when edf is zero the term is effectively dropped from the model. The test statistic reported is *χ*^2^ for survival and flowering and *F* for log(size_t+1_).

**Figure 3:**
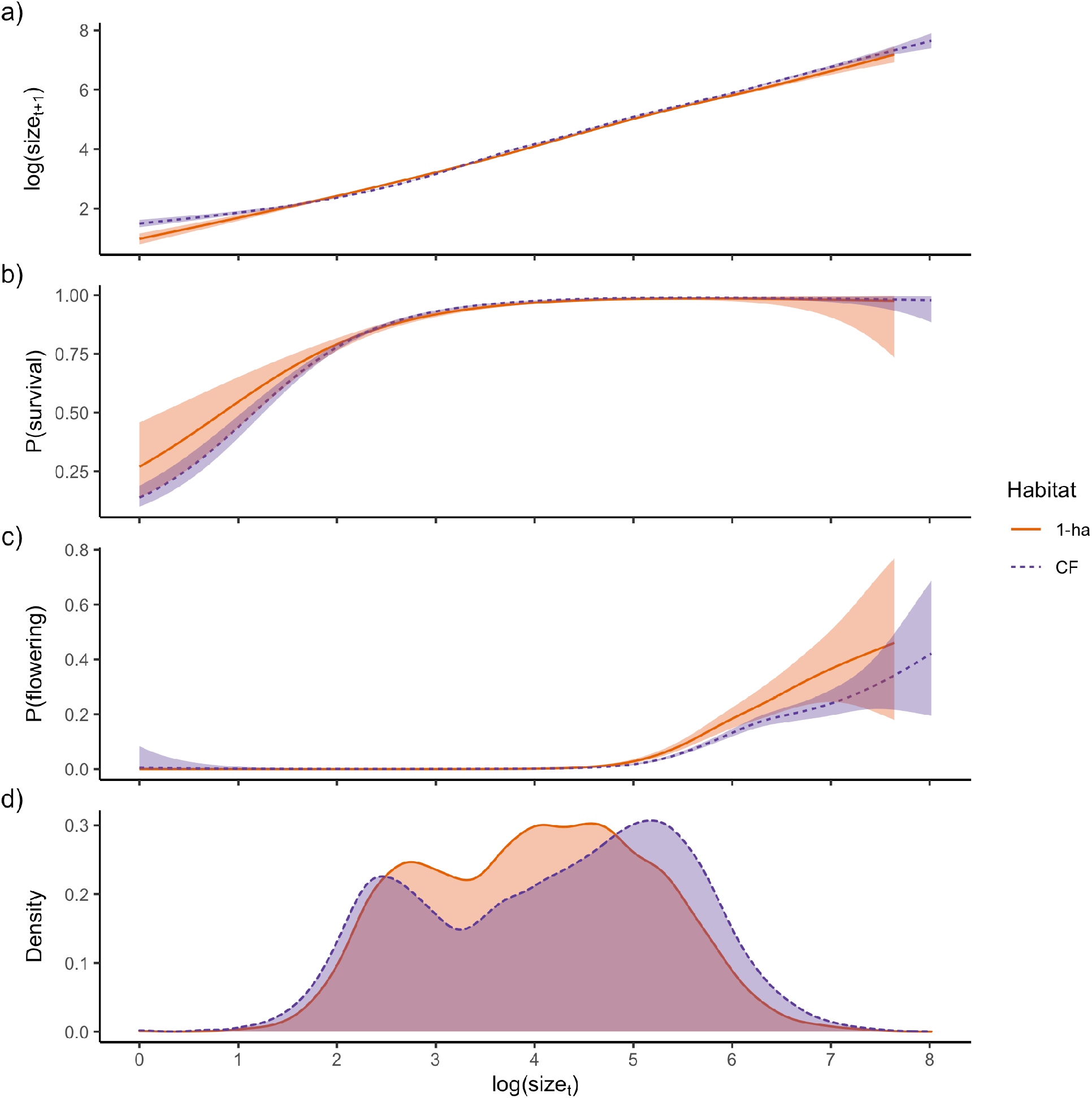
Marginal effect of plant size in the previous census on (a) survival, (b) log(size), and (c) flowering probability. Plots a–c were created by evaluating the smooth functions of *log*(*size_t_*) at observed values, adding the model intercepts, and back-transforming (when appropriate) to the response scale (e.g. probability for a and c). The bands depicting the 95% confidence interval include uncertainty in the intercept and uncertainty due to smoothness selection; the smooths for 1-ha fragments and continuous forest are fit in separate models. (d) Plant size distribution by habitat type (solid line = 1-ha fragments, dashed line = Continuous Forest). The curves in a–c are shown with raw data superimposed in Figure S3.

#### Size

Plants in continuous forest had an average of 2.9 shoots (± 1.8 SD) and were on average 39.8 cm tall (± 26.2 SD). Plants in 1-ha fragments had on average 14% fewer shoots (2.5 ± 1.4 SD) and were 10.5% shorter (35.6 cm ± 23.8 SD). Because our proxy for plant size was the product of these two metrics, plants in continuous forest were on average 32% larger than those in forest fragments (144 ± 176 SD vs. 109 ± 139 SD, respectively), with fragments having proportionately fewer large plants (Figure 3d). Plants were significantly larger in continuous forest on average (Table 1), and the disparity in plant size—which was most pronounced in the initial years of our surveys—diminished over time.

Mean plant size dropped dramatically in 2003 in both habitat types (negative fold-change in Figure 1a), corresponding with a severe drought during the February census (SPEI = - 1.39) (Figure 1d). As with survival, size in year t was a significant predictor of size in year t+1 (Table 2). Although plants were significantly larger in continuous forest compared to fragments (Table 1), the effect of size in year t on size in year t+1 is nearly identical in the two habitats (Figure 3a).

#### Flowering

The overall proportion of plants that flowered was very low, but the proportion of plants flowering was 31% higher in continuous forest than 1-ha fragments (4.6% vs. 3.5%, respectively). The intercepts for the continuous forest and 1 ha fragment models were significantly different, indicating a main effect of habitat on flowering probability (Table 1). The observed disparity in proportion of flowering plants was largely due to the fact that flowering is also significantly size-dependent (Table 2), with the probability of flowering increasing dramatically once plants reached the threshold size of about 148 (i.e., log(size) > 5 in Figure 3c). Despite the flowering probability of the largest plants being greater in fragments than continuous forest, populations in fragments had proportionately fewer plants above the reproductive size threshold (Figure 3d). The most striking differences between habitat types coincided with relatively extreme weather conditions. In 2003, during the onset of a severe drought, the proportion of potentially reproductive plants (i.e. plants above the size-threshold for reproduction) that actually flowered was 28% in continuous forest vs. 17% in 1-ha fragments. In 2006, following a wet spring and winter (SPEI > 1), the trend was reversed: only 14% of these plants flowered in continuous forest vs. 26% in 1-ha fragments (Figure 1c). Note that plots in 1-ha fragments generally have fewer and smaller plants than those in continuous forest, so total numbers of flowering plants was always lower in fragments.

### Delayed effects of drought on demographic vital rates

Drought history had a significant effect on the survival, growth, and flowering of plants in both habitats (Table 2). Comparing the respective crossbasis surfaces, however, reveals that the specific climatic drivers, their timing, and their impact on individual vital rates all differed among habitats.

#### Survival

For 1-ha fragments, there was a significant effect on survival of SPEI in the preceding 0–16 months. The highest survival was near SPEI of 0, with mortality increasing as conditions became either drier or wetter (i.e., as SPEI values became increasingly negative or positive, respectively; Figure 4b). Additionally, a positive, lagged effect of high SPEI at 32–36 months lag reached statistical significance. There was less of a delayed effect of SPEI on survival in continuous forest with only the preceding 0–4 months and 32–36 months reaching statistical significance (Figure 4a). The short-term effects of SPEI on survival in continuous forest were also unidirectional—the probability of survival was highest in wet conditions and declined, albeit only slightly, with increasingly negative values of SPEI (i.e., as droughts became more severe; Figure 4a). Plants in both habitat types showed an increase in survival probability with very high SPEI values (i.e., extremely high precipitation) at a lag time of 32–36 months. It should be noted, however, that only the first transition year of census data (1998–1999) met these conditions. We compared the effects of SPEI history in continuous forest and fragments by subtracting the fitted values in Figure 4b from Figure 4a to produce Figure 4c. This shows that in average conditions (SPEI = 0), there is little difference in survival probability between continuous forest and forest fragments (Figure 4c). However, under extreme conditions, survival probability is up to 0.042 higher in continuous forest than fragments.

**Figure 4:**
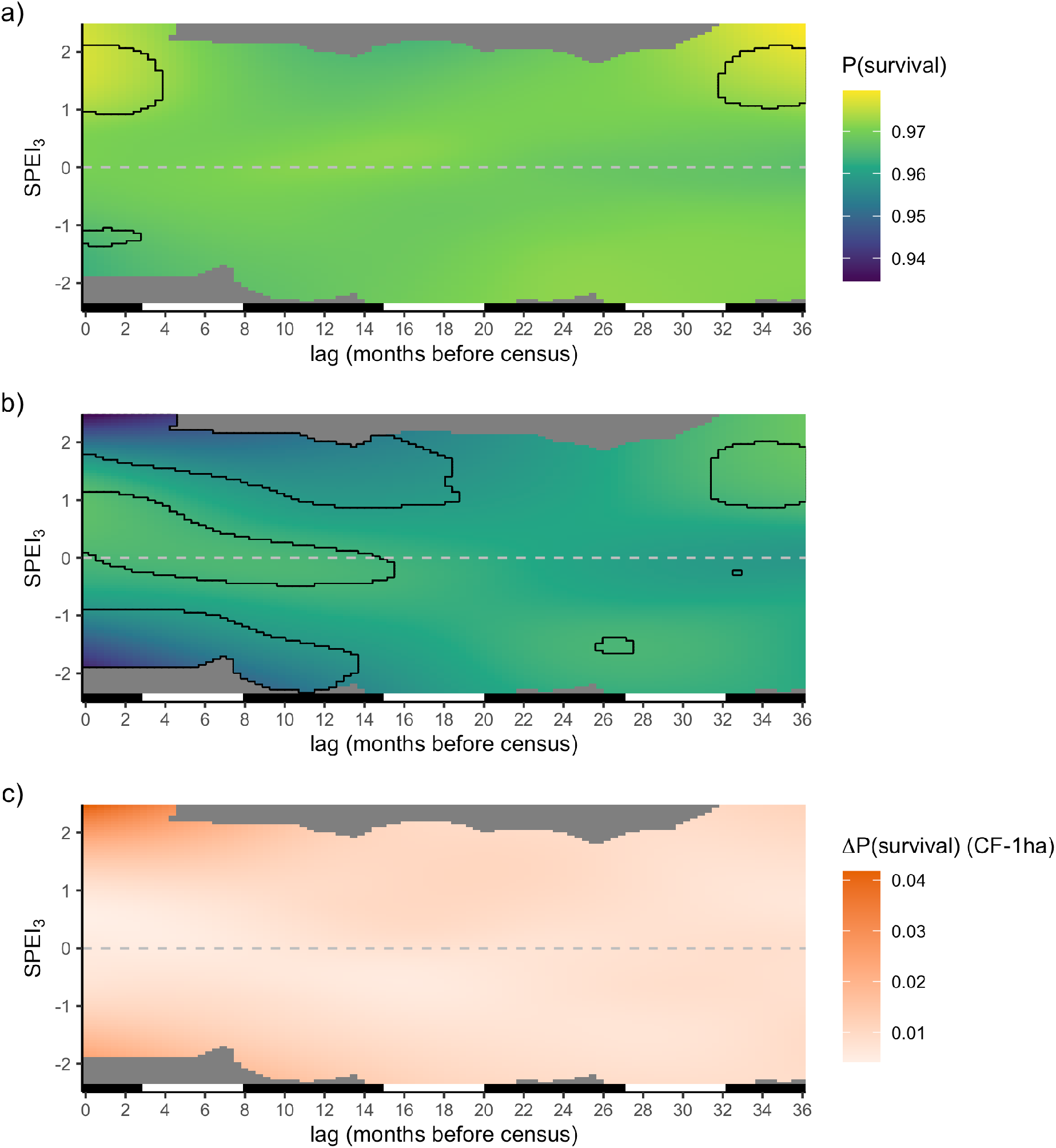
Marginal effect of lagged SPEI on *H. acuminata* survival in (a) continuous forest, (b) 1-ha fragments, (c) and the difference between the two. Outlines show combinations of SPEI and lag time that have a significant effect on survival, defined as areas where the 95% confidence interval around the response does not overlap the intercept. The bar on the bottom of each panel indicates wet seasons (black, November–May) and dry seasons (white, June–October). For a and b, the model intercepts were added to the evaluated crossbasis smooths and values were back-transformed to the response scale (i.e. probabilities). Areas of the fitted smooth far from observed values (i.e. combinations of lag time and SPEI) are shown in grey.

#### Size

The effects of drought history on trends in plant size were generally similar for continuous forest and fragments. Under all conditions, plant size is greater in continuous forest (Figure 5c). Decreasing SPEI at lags of 8–11 months (i.e., the end of the preceding year’s wet season) led to increased growth in both habitats. In continuous forest, but not 1- ha fragments, SPEI at lags of 22–25 months had a significant effect on plant size in the opposite direction, with wet conditions at that lag time resulting in larger predicted sizes. At longer lags of 26–36 months positive effects of drought on plant size are predicted for both continuous forest and 1-ha fragments by our models, although the area of SPEI–lag space that reaches statistical significance is larger for continuous forest.

**Figure 5:**
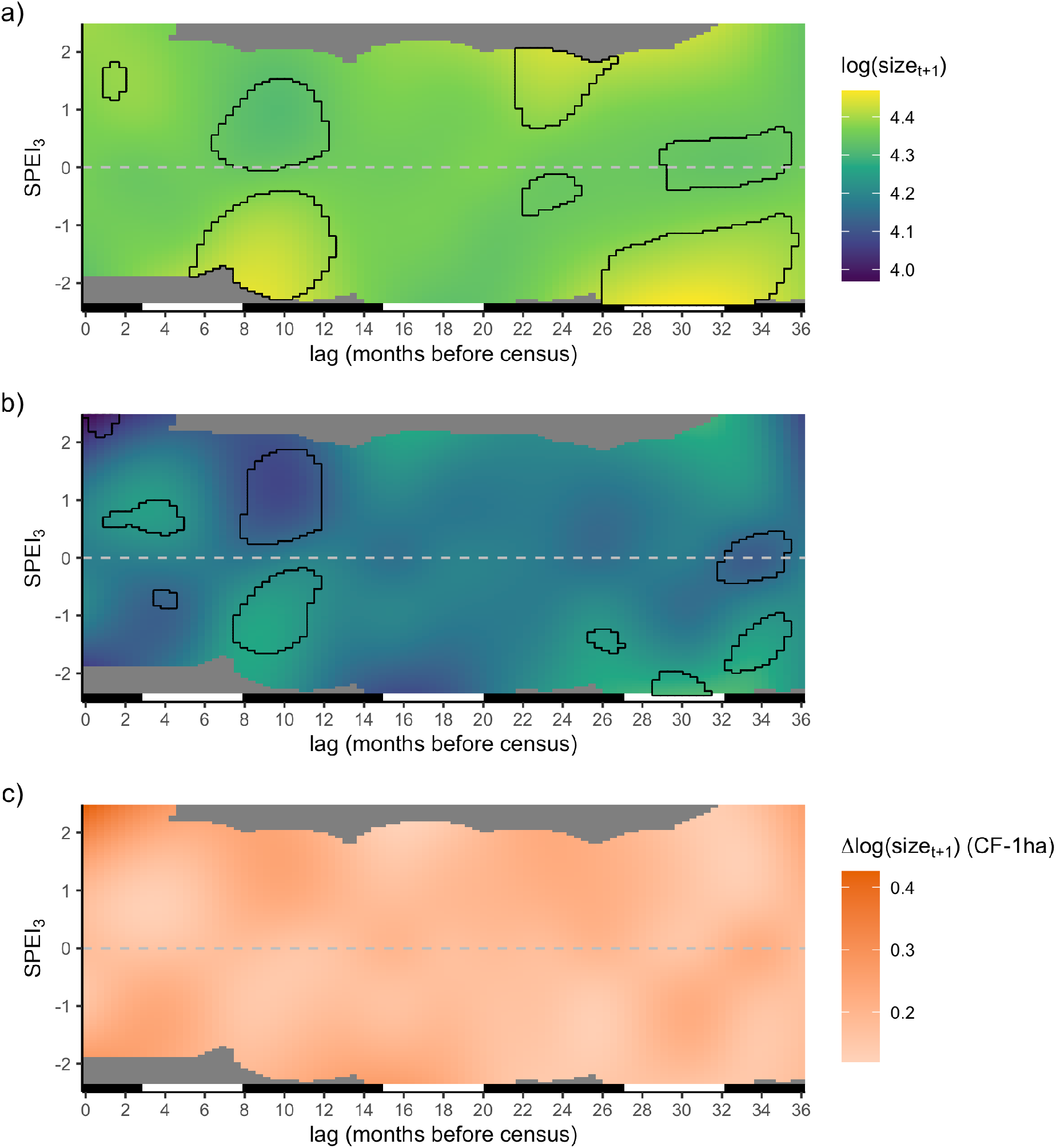
Marginal effect of lagged SPEI on *H. acuminata* size in (a) continuous forest, (b) 1-ha fragments, (c) and the difference between the two. Outlines show combinations of SPEI and lag time that have a significant effect on plant size, defined as areas where the 95% confidence interval around the response does not overlap the intercept. The bar on the bottom of each panel indicates wet seasons (black, November–May) and dry seasons (white, June–October). For a and b, the model intercepts were added to the evaluated crossbasis smooths. Areas of the fitted smooth far from observed values (i.e. combinations of lag time and SPEI) are shown in grey.

#### Flowering

Overall, the probability of flowering was significantly higher in continuous forest than in 1-ha fragments (Figure 6, Table 1). Recent and past SPEI had less of an effect on flowering probability in 1-ha fragments as indicated by the narrower range of the evaluated smooth (Figure 6b). This led to some important inter-habitat differences in plant responses to prior droughts. In continuous forests, recent drought (i.e., at lag = 0–2 with SPEI < -1) and droughts two dry seasons prior (lags 15–20) increased the probability of flowering. The shape of the crossbasis smooth for 1-ha fragments suggested that moderate drought (-1.5 <SPEI< -1) in the previous 0–18 months and at lags of 28–36 months significantly increased flowering probability slightly (the highest probability estimated is 0.004; Figure 6b). The effects of drought on flowering probability appeared stronger in continuous forest compared to 1-ha fragments (Figure 6c). We found no evidence for a cost of reproduction: in both forest and fragments, plants that had flowered in the previous year were significantly more likely to be larger and flower again. Finally, with the exception of the model for survival in 1-ha fragments, the random effect of plot was significant, indicating that vital rates varied among plots (Table 2).

**Figure 6:**
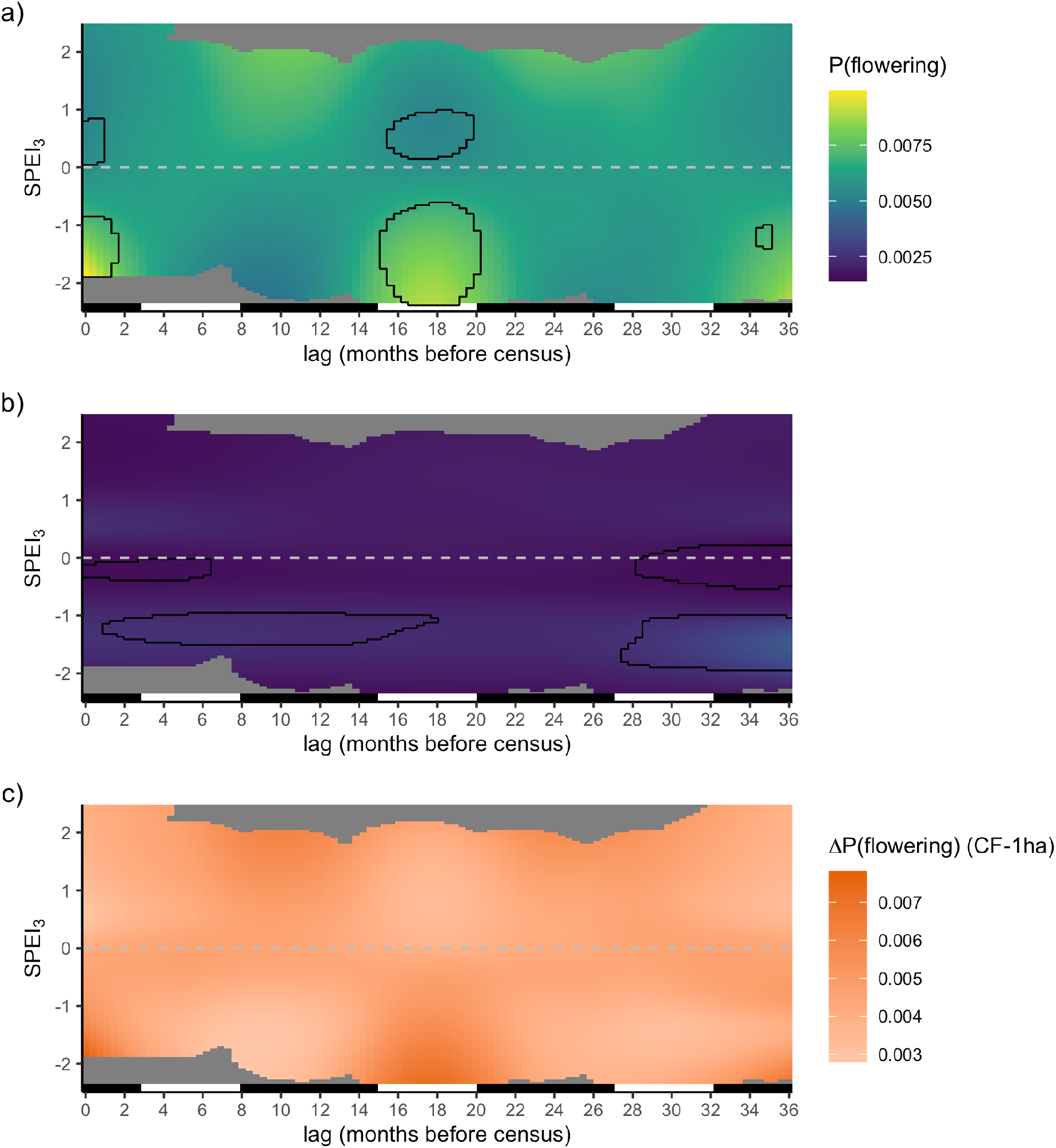
Marginal effect of lagged SPEI on *H. acuminata* flowering probability in (a) continuous forest, (b) 1-ha fragments, (c) and the difference between the two. Outlines show combinations of SPEI and lag time that have a significant effect on probability of flowering, defined as areas where the 95% confidence interval around the response does not overlap the intercept. The bar on the bottom of each panel indicates wet seasons (black, November–May) and dry seasons (white, June–October). For a and b, the model intercepts were added to the evaluated crossbasis smooths and values were back-transformed to the response scale (i.e. probabilities). Areas of the fitted smooth far from observed values (i.e. combinations of lag time and SPEI) are shown in grey.

## Discussion

Understanding how landscape structure and abiotic conditions act to influence population dynamics is central to many conceptual frameworks for studying and conserving fragmented landscapes (Didham et al., 2012; Driscoll et al., 2013). Our results support the emerging consensus that the effects of climatic extremes on demographic vital rates can be delayed for months or even years (Evers et al., 2021; Teller et al., 2016; Tenhumberg et al., 2018). We also found different magnitudes, directions, and lag times of climate effects in fragments and continuous forests. This suggests that the hypothesized synergies between climate and fragmentation on population dynamics (Laurance & Williamson, 2001; Opdam & Wascher, 2004; Selwood et al., 2015) may be important in this system in a way far more complex than previously thought.

### Temporal variation in demographic responses to forest fragmentation

Many studies investigating the biological consequences of habitat fragmentation on plant growth, survival, and reproduction comprise short-term (≤3 year) experiments and observations. Our results underscore the difficulty in extrapolating long-term trends from such short-term studies, particularly when studying long-lived organisms or when the responses of interest can vary with size or age. For instance, one would have reached a very different conclusion regarding the effect of fragmentation on annual survival if the study windows were 1999–2002 (i.e., higher survival in continuous forest), 2002–2005 (i.e., higher survival in fragments), or 2004–2007 (i.e, no clear effect of fragmentation) (Figure 1b). It is only when evaluating over longer time windows that it becomes apparent mortality is elevated in fragments relative to continuous forest (Figure 2), and that the observed interannual variation is largely driven by dynamic patterns of recruitment (Bruna, 2002) coupled with low mortality for plants beyond the smallest size classes (Bruna, 2003).

Similarly, conclusions regarding the effects of fragmentation on flowering—which is also both rare and size-dependent (Brooks et al., 2019)—would also differ based on the year in which they were investigated. This could lead to erroneous extrapolations regarding the effects of fragmentation on reproductive mutualists or population genetic structure (Côrtes et al., 2013; Uriarte et al., 2010; Uriarte et al., 2011). Conclusions based on short-term observations of temporally variable vital rates could lead to conservation and management practices that are ineffective or even counterproductive, especially when when failing to consider how the consequences of this variation might be modulated by organismal life history (Morris et al., 2008).

It is important to emphasize, however, that the range of the estimated response to SPEI for survival and growth was greater in fragments compared with continuous forest (Figures 4, 5). This suggests that extremes in SPEI may be more detrimental in forest fragments compared with continuous forest. While intact forest and its canopy buffer populations from climatic extremes, populations in fragments—especially near edges with high contrast matrix—likely lack this protection (Didham & Lawton, 1999; Ewers & Banks-Leite, 2013). We suggest it is these climate extremes, rather than trends in average temperature, precipitation, or SPEI (Laurance et al., 2014), that that are the causal mechanism underlying the observed reduced plant survival and size in forest fragments.

### Delayed effects of climate on demographic vital rates

Climate anomalies are known to have immediate effects on the growth, survival, or reproduction of plants (Esteban et al., 2021; Wright & Calderon, 2006), including *Heliconia* (Stiles, 1975; Westerband et al., 2017) and other tropical herbs (Wright, 1992). These effects can be complex or even contradictory—mild droughts can increase the growth rates of tropical trees and seedling survival, perhaps due to reductions in cloud cover and concomitant increases in solar radiation (Alfaro-Sánchez et al., 2017; Condit et al., 2004; Huete et al., 2006; Jones et al., 2014; Uriarte et al., 2018), but in severe drought years growth can be extremely low and mortality can be sharply elevated (Connell & Green, 2000; Edwards & Krockenberger, 2006; Engelbrecht et al., 2002). There is also evidence that the effects can persist for multiple years (Phillips et al., 2010), such as a boom in drought-year fruit production followed by severe post-drought “famine” (Pau et al., 2013; Wright et al., 1999).

Despite these insights, models of plant population dynamics rarely include the effects of environmental drivers [but see Williams et al. (2015); Tenhumberg et al. (2018); Molowny-Horas et al. (2017)). This has largely been due to the challenge (both ecologically and statistically) of detecting any demographic responses to climatic extremes that are delayed for multiple growing seasons. To address this, researchers have begun to use a number of statistical methods that test for time lags in demographic responses without *a priori* assumptions about the influence of any particular climate window (Evers et al., 2021; Ogle et al., 2015; Teller et al., 2016; Tenhumberg et al., 2018). Our results are consistent with this emerging literature—that the effects of precipitation extremes on the demography of *Heliconia acuminata* could be delayed for up to 3 growing seasons. Additionally, our method allowed us to capture significant non-linear responses to SPEI at different lag times, which appear to differ in shape between habitats.

While it appears that delayed effects of climate on demographic vital rates may be ubiquitous (Evers et al., 2021), the extent to which they vary spatially or with habitat remains an open question. Our results suggest that they may, with habitat-specific differences in how environmental conditions influenced future vital rates. For example, extreme values of SPEI—both positive (unusually high precipitation) and negative (drought conditions)—led to declines in the probability of individual survival in forest fragments while extreme wet conditions significantly increased survival in continuous forest. Similarly, the marginal effects of SPEI on plant size were greater in fragments, suggesting a more pronounced effect of extreme climates in fragments. In contrast, variation in SPEI corresponded to a greater range of flowering probabilities in continuous forest than fragments. These results should be interpreted with some caution, however, as the relatively low number of plants in fragments that are above the threshold-size for flowering could limit the power to detect delayed effects.

Interestingly, we found significant detrimental effects of unusually wet conditions at some lag times for all vital rates except survival in continuous forest and flowering probability in 1-ha fragments. While most studies investigating how precipitation extremes influence tropical plants have focused on droughts (Lewis et al., 2011; Phillips et al., 2009; Williamson et al., 2000), more recent work has shown that unusually wet conditions can also be detrimental (Esteban et al., 2021). For instance, the increased cloud cover associated with elevated precipitation could lead to reduced photosynthetic activity and growth; plants in the resulting saturated soils could also have lower growth and elevated mortality (Parent et al., 2008). The wind storms accompanying extreme precipitation events (Espírito-Santo et al., 2010; Negrón-Juárez et al., 2018) could increase the likelihood of tree-, branch-, and litter-fall, all of which are sources of mortality for understory plants (Scariot, 2000; Ssali et al., 2019). Finally, cool temperatures associated with unusually wet conditions may decrease flowering (Pau et al., 2013).

There are several, non-mutually-exclusive explanations for delayed effects of SPEI on demography. The first is that the physiological processes underlying vital rates might be initiated long before they are demographically apparent (Evers et al., 2021), and hence be shaped by climatic events at any point in that physiological window. For example, the flowering shoots of *Heliconia chartacea* begin to develop 6–10 months prior to the appearance of inflorescences (Criley & Lekawatana, 1994). Adverse conditions during the 6 months following initiation, rather than the months when inflorescences are starting to expand, leads to the aborted production of flowering shoots. Our results showed delayed increases in flowering probability after drought. Drought conditions could be favorable for *H. accuminata* flowering due to increased temperatures or decreased cloudiness associated with droughts (Pau et al., 2013), and the effects could be delayed due to the development time of inflorescences.

Demographic responses will also be delayed if abiotic stress causes plants to invest in belowground rhizomes (*sensu* Pumisutapon et al., 2012). The carbohydrates stored in rhizomes allow *Heliconia* to regenerate aboveground biomass following damage (Rundel et al., 1998) and protect the buds that give rise to new shoots from stressful conditions (Klimešová et al., 2018). This may be why drought led to delayed increases in growth—by shedding shoots and leaves and investing in rhizomes, plants could be generating proportionately more buds with which to regenerate when conditions improve. This is consistent with the results of prior experiments in which *H. acuminata* plants transplanted into hotter, drier fragments lost and then recovered far more leaf area than control plants (Bruna et al., 2002).

Third, it may be that the delayed demographic effects we observed are indirectly mediated by the effect of SPEI on other species rather than the direct effects on individual physiology (Evers et al., 2021). For example, topical trees may not die until three or more years after a drought (Criley & Lekawatana, 1994). When they finally do, the resulting leaf drop (Janssen et al., 2021) and treefalls allow for light penetration to the forest understory (Canham et al., 1990; Leitold et al., 2018), triggering a boom in the growth and flowering of understory plants (Bruna & Oli, 2005). Similar delayed changes in the local environment could also influence the foraging behavior of a plant’s pollinators (Bruna et al., 2004; Stouffer & Bierregaard, 1996), seed dispersers (Uriarte et al., 2011), or herbivores (Scott et al., 2021). While more work is needed to explain why the (delayed) effects of SPEI on *H. acuminata* survival and growth are greater in fragments than forest interiors, one hypothesis, motivated by recent intriguing results from other systems (Sapsford et al., 2017), is that the greater litterfall on edges (Vasconcelos & Luizão, 2004) may be altering the abundance of pathogens or mycorrhizae.

Finally it is important to clarify the distinction between “physiological delays” and “observational delays.” For example, a Fall drought could reduce cold tolerance and therefore overwinter survival. Alternatively the drought could kill plants immediately. If mortality is recorded in Spring, then mortality would appear as a delayed effect of the drought in both hypothetical cases, but only the first case is a truly delayed physiological response to drought. The apparent delayed response in the second case is due to the timing of the response in relation to the census date. In our analysis, this potential explanation of “observational delays” applies only to plant size and survival, as the flowering season coincided with the yearly census. It also only applies to lags of up to 12 months since mortality, size, and flowering are recorded yearly. This possibility is not unique to our study, rather it is a consequence of conducting demographic censuses on an annual scale while the climate is quantified monthly or seasonally. For some demographic models and follow-up experiments, knowing the precise timing of mortality might be critical; for others less so. To disentangle possible mechanisms for observed delayed effects, it may be necessary to conduct demographic surveys at the same temporal scale at which climate is aggregated.

## Conclusions & Future Directions

Over 24 million ha of the Brazilian Amazon have been cleared in the last two decades (Silva Junior et al., 2021), resulting in their extensive fragmentation (Broadbent et al., 2008). Climate models predict a future of extremes for these forests—increases in the frequency and geographic extent of droughts, but also increases in the frequency and area affected by periods of unusual wetness (Duffy et al., 2015). Our results support the hypothesis that populations in Amazonian forest fragments could be more susceptible to the effects of changing climate than those in continuous forest (Laurance et al., 2014). However, they also indicate that the demographic responses to climate change of populations in fragmented landscapes may be far more complex than previously appreciated. Multi-factorial, multi-season experiments (*sensu* Aguirre et al., 2021; Bruna & Ribeiro, 2005; Markewitz et al., 2010; Westerband et al., 2017), ideally manipulating multiple combinations of climatic variables (Mundim & Bruna, 2016), are needed to determine how and why habitat-specific differences in environmental conditions interact to delay the demographic responses of plants to climatic variability. Also needed are statistical tools that can test for synergistic effects of fragmentation and climate in vital rates, as those currently available do not allow for including interaction terms. This also limits the ability to include size by climate interactions in a DLNM; although plant responses to both fragmentation and climatic extremes can be size-specific (Bruna & Oli, 2005; Schwartz et al., 2019). The ability to identify size-specific lagged responses may be especially complicated given size is rarely measured at the same time scale (e.g. monthly) as climate drivers.

Recent research, including our own presented here, shows that lagged effects of climate drivers on demography may be the norm (Evers et al., 2021). As such, we suggest that all demographers investigate the possibility of lagged environmental effects, ideally using methods that do not require an *a priori* choice of lag times such as DLNMs, automated algorithms for choosing critical weather windows (van de Pol et al., 2016), or Bayesian methods that incorporate “ecological memory” (Ogle et al., 2015).

Finally, any analytical approach for assessing lagged effects on demography requires long-term data and/or data collected from many sites with independent weather (but similar climate and site conditions) (Evers et al., 2021; Tenhumberg et al., 2018). Teller et al. (2016) used a simulation study to show that detecting lagged effects required 20–25 years of data, although this depended on effect size and the range of climate extremes in the data. Work in progress from Compagnoni et al. (2021) showed that spatial replication may alleviate this requirement, and in fact in our study we were able to detect significant lagged effects with only 10 years of data and multiple plots experiencing slightly different weather. Unfortunately, long-term data monitoring the entire life-cycle of tropical taxa are rare, and those doing so in fragmented landscapes are virtually nonexistent (Bruna & Ribeiro, 2005). Without investing in collecting such data, generalizations regarding the demographic consequences of climate change in these species rich and increasingly fragmented habitats will continue to prove elusive. More generally, however, researchers need to consider how delayed responses to climate could influence the interpretation of data in studies where the organisms lifespan exceeds the study’s duration.

## Acknowledgments

We thank Collin Edwards, Andrew Mercadante, Ellie McDaniel and 3 anonymous reviewers for helpful discussions and comments on the manuscript. We also thank the technicians and students who helped conduct the *Heliconia* censuses and the BDFFP and INPA for logistical support. Financial support was provided by the U.S. National Science Foundation (awards DEB-0614339, DEB-0614149, INT 98-06351, and DEB-1948607). This is publication No. in the BDFFP Technical Series. The authors delcare no conflicts of interest.

## Data Availability Statement

Data used in this study are available at the Dryad Digital Repository [links included upon acceptance]; R code for analyses and visualizations are archived with Zenodo at https://doi.org/10.5281/zenodo.5364591.

