## Appendix A for "Delayed effects of climate on vital rates lead to demographic divergence in Amazonian forest fragments"

**Reproducibility** **15**

#### Survival

##### Forest Fragments

###### Diagnostics

```
s_1ha_qres <- qresid(s_1ha)
par(mfrow = c(2,2))
#histogram
plot(density(s_1ha_qres))
#QQ plot
```

```
qqnorm(s_1ha_qres); qqline(s_1ha_qres)
#Fitted vs. residuals--binned residuals plot
binnedplot(fitted(s_1ha), residuals(s_1ha, type = "response"))
```

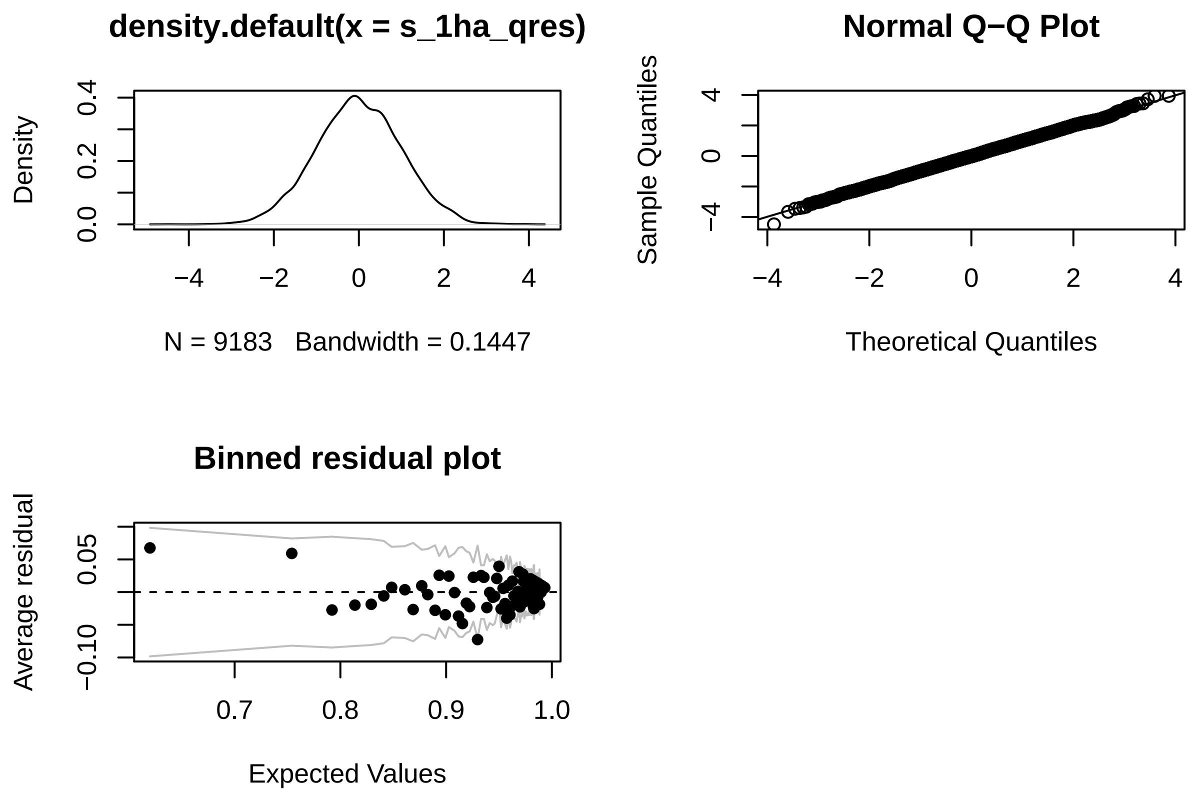

##### Basis dimensions

```
k.check(s_1ha)
```

```
##                       k'          edf   k-index p-value
## s(log_size_prev)    9 2.842178e+00 0.9304969   0.845
## s(spei_history,L) 90 1.077927e+01        NA      NA
## s(plot)             4 1.627947e-04        NA      NA
```

`k` is adequate for `log_size_prev`. Unfortunately `k.check()` doesn't work for data with matrices, so I'll use an alternative approach detailed in `?mgcv::choose.k` for the crossbasis smooth. Briefly, it involves looking for pattern in residuals by extracting residuals and fitting a GAM to them with the smooth of interest with an increased value for `k`. If there is no unexplained pattern in residuals, that smooth should have an edf near zero.

```
check_res_edf <- function(model) {
  res <- residuals(model)
  mgcv::gam(res ~ te(spei_history, L, k = c(20, 35), bs = "cs"),
```

```r
      gamma = 1.4,
      data = model.frame(model)) %>%
    gratia::edf() %>%
    dplyr::mutate(edf = round(edf, 1))
}

check_res_edf(s_1ha)
```

```
## # A tibble: 1 x 2
##   smooth               edf
##   <chr>              <dbl>
## 1 te(spei_history,L)     0
```

Adequate k for crossbasis.

#### Continuous Forest

##### Diagnostics

```r
s_cf_qres <- qresid(s_cf)
par(mfrow = c(2,2))
#histogram
plot(density(s_cf_qres))
#QQ plot
qqnorm(s_cf_qres); qqline(s_cf_qres)
#Fitted vs. residuals--binned residuals plot
binnedplot(fitted(s_cf), residuals(s_cf, type = "response"))
```

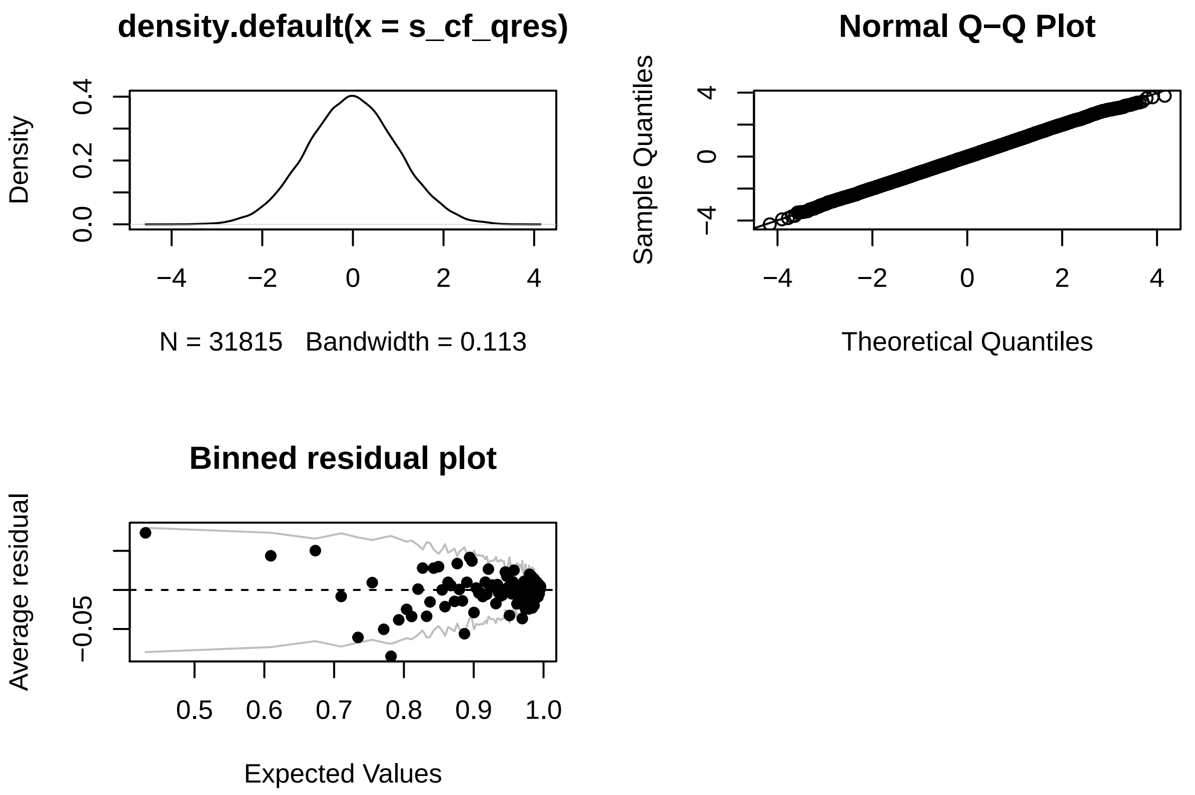

**Basis dimensions**

```
k.check(s_cf)
```

```
##                      k'       edf  k-index p-value
## s(log_size_prev)      9  3.475276 0.957276    0.96
## s(spei_history,L)   210 12.871869       NA      NA
## s(plot)               6  4.361255       NA      NA
```

Adequate knots for `log_size_prev`

```
check_res_edf(s_cf)
```

```
## # A tibble: 1 x 2
##   smooth                edf
##   <chr>               <dbl>
## 1 te(spei_history,L)      0
```

Adequate knots for crossbasis smooth.

##### Effect of sample size

Here I use a random sub-sample to check that differences between continuous forest and fragments are not purely due to sample size differences, particularly differences in the complexity and shape of the crossbasis smooth.

```
summary(s_1ha)$edf[2]
```

```
## [1] 10.77927
```

```
summary(s_cf)$edf[2]
```

```
## [1] 12.87187
```

```
summary(s_cf_sub)$edf[2]
```

```
## [1] 1.80754
```

The crossbasis is smoother, but shows a similar pattern.

```
draw(s_cf, select = "s(spei_history,L)", n_contour = 5)
```

```
## Warning: Removed 939 rows containing non-finite values (stat_contour).
```

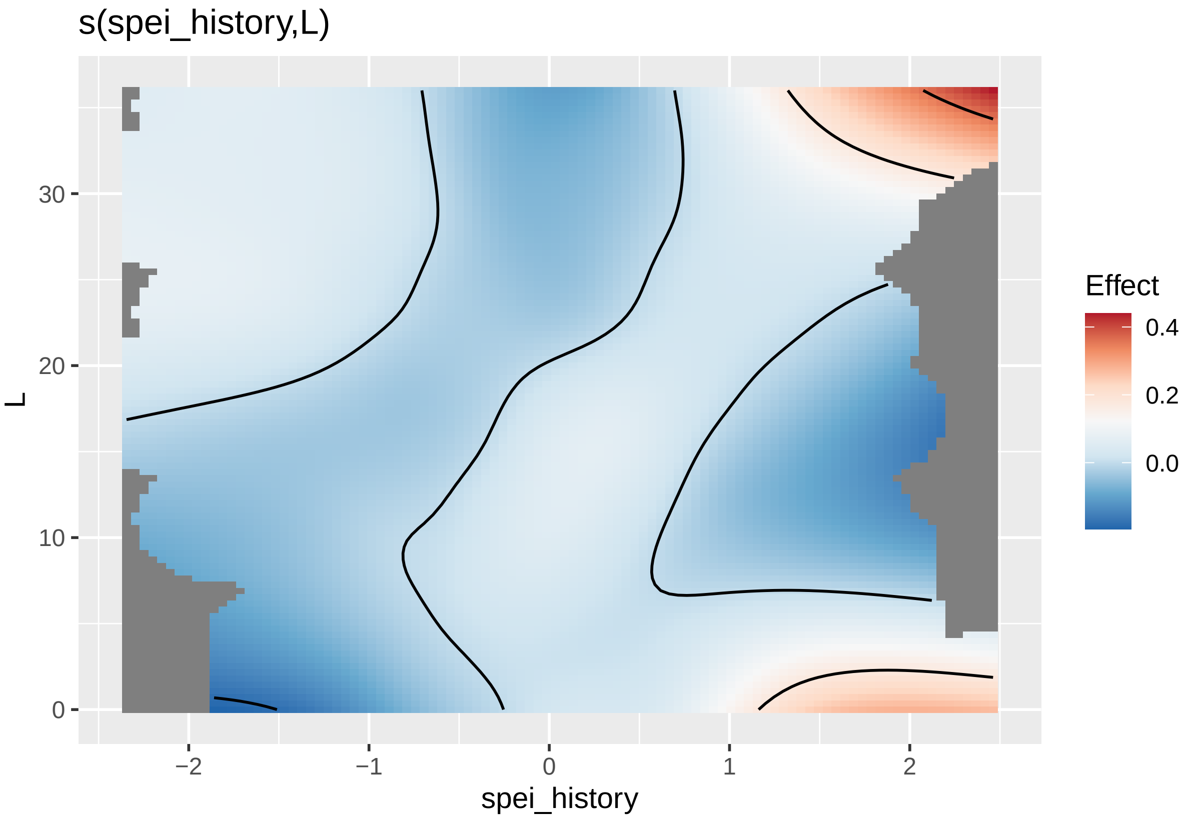

```r
draw(s_cf_sub, select = "s(spei_history,L)", n_contour = 5)
```

```
## Warning: Removed 910 rows containing non-finite values (stat_contour).
```

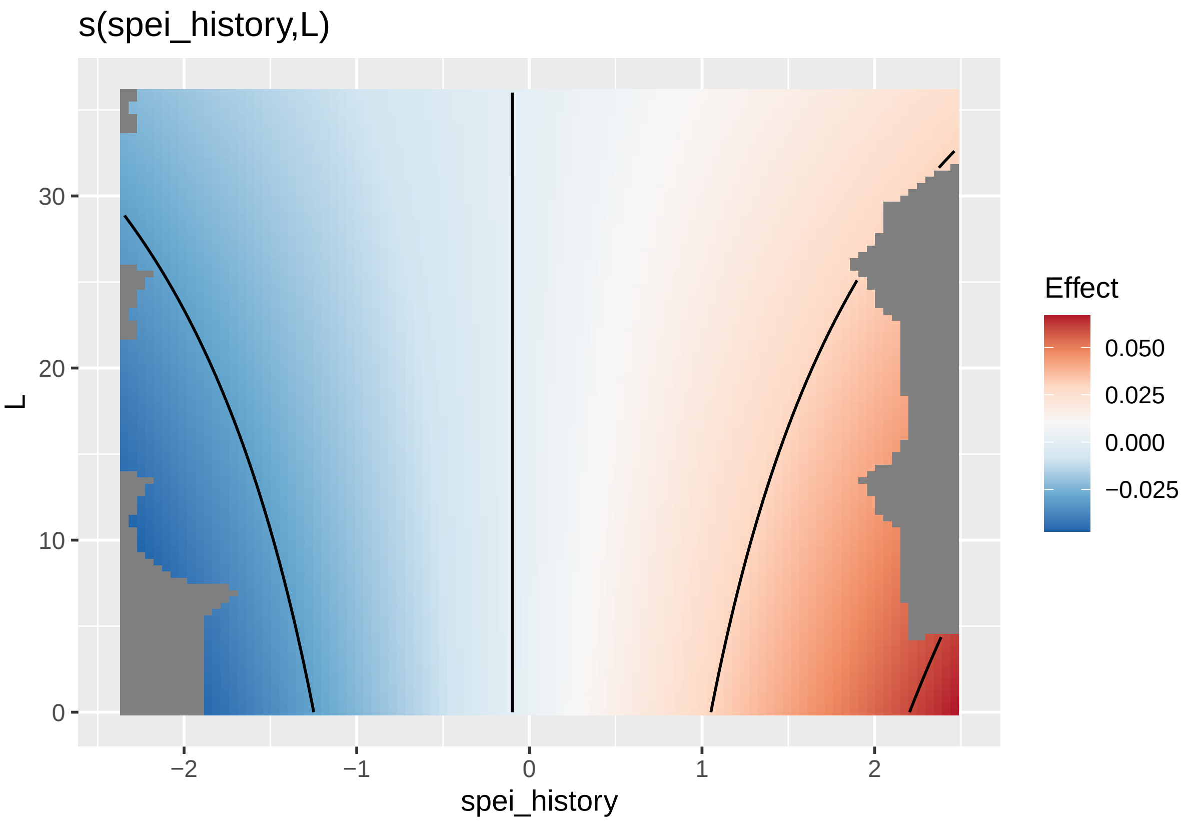

### Growth

#### Fragments

##### Diagnostics

```r
appraise(g_1ha)
```

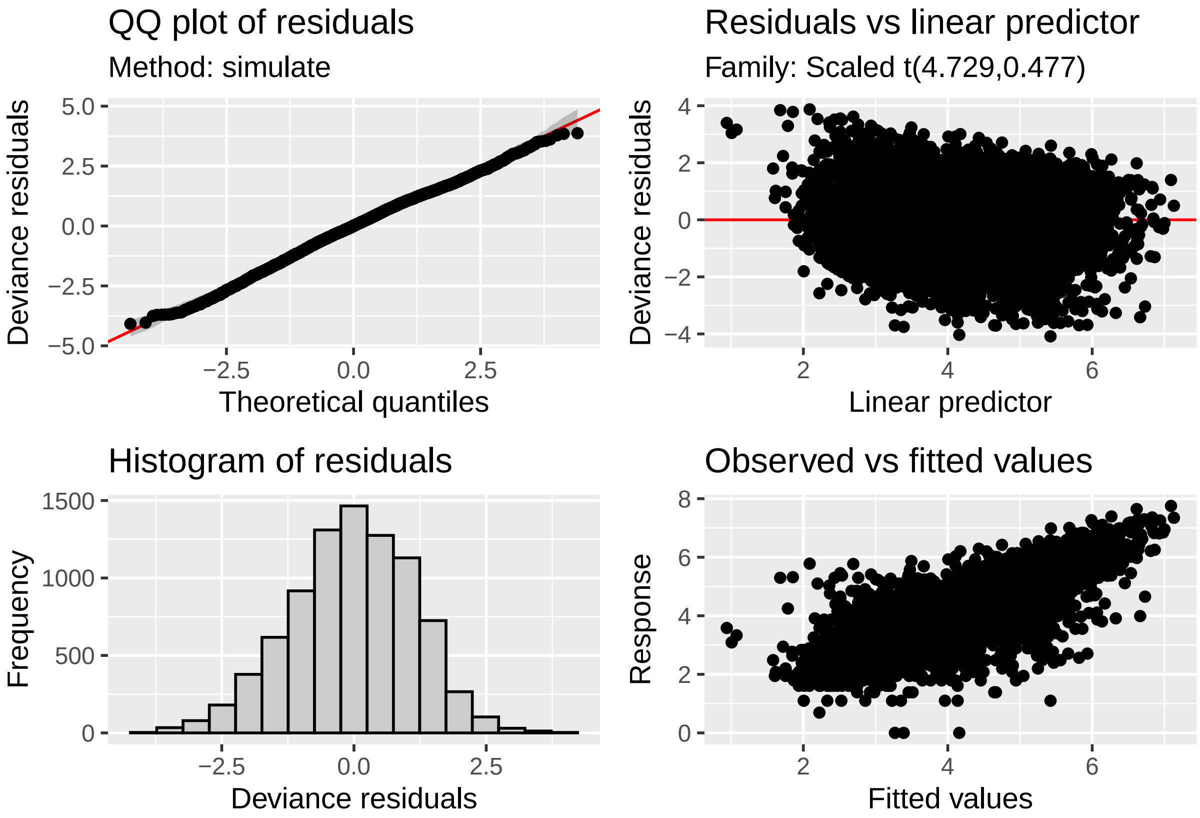

**Basis dimensions**

```
k.check(g_1ha)
```

```
##                     k'       edf   k-index p-value
## s(log_size_prev)    9  4.193965 0.9717753    0.02
## s(spei_history,L)  60 18.181554        NA      NA
## s(plot)             4  2.826248        NA      NA
```

Adequate k for `log_size_prev`.

```
check_res_edf(g_1ha)
```

```
## # A tibble: 1 x 2
##   smooth              edf
##   <chr>             <dbl>
## 1 te(spei_history,L)     0
```

Adequate k for crossbasis.

#### Continuous Forest

##### Diagnostics

```
gratia::appraise(g_cf)
```

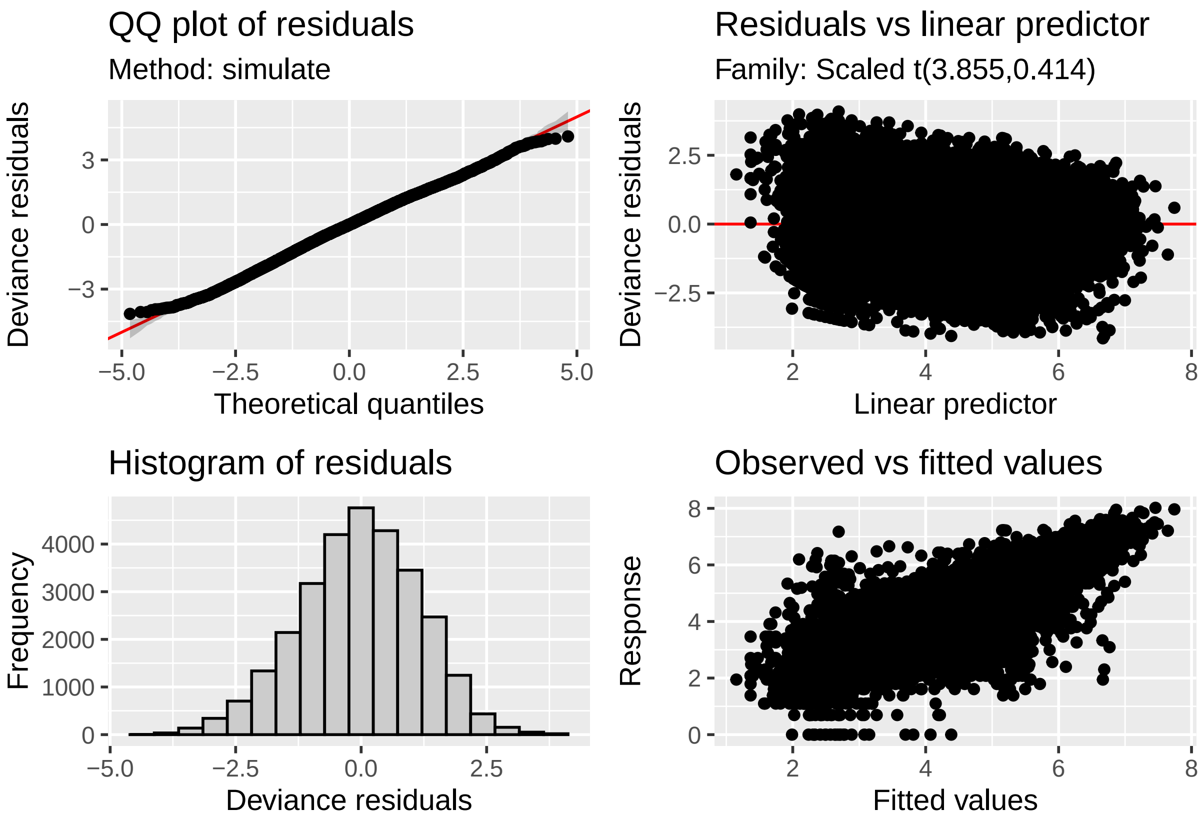

##### Basis dimensions

```
k.check(g_cf)
```

```
##                      k'       edf   k-index p-value
## s(log_size_prev)   24  9.304260 0.9744131  0.0225
## s(spei_history,L)  60 15.398732        NA      NA
## s(plot)             6  4.016572        NA      NA
```

Adequate k for `log_size_prev`

```
check_res_edf(g_cf)
```

```
## # A tibble: 1 x 2
##   smooth                edf
##   <chr>               <dbl>
## 1 te(spei_history,L)      0
```

Adequate k for crossbasis smooth.

##### Effect of sample size

To check that differences are not purely due to sample size differences, particularly that lower edf in continuous forests is due to higher sample size.

```
summary(g_1ha)$edf[2]
```

```
## [1] 18.18155
```

```
summary(g_cf)$edf[2]
```

```
## [1] 15.39873
```

```
summary(g_cf_sub)$edf[2]
```

```
## [1] 14.09059
```

edf is similar for the sub-sample

```
draw(g_cf, select = "s(spei_history,L)", n_contour = 5)
```

```
## Warning: Removed 968 rows containing non-finite values (stat_contour).
```

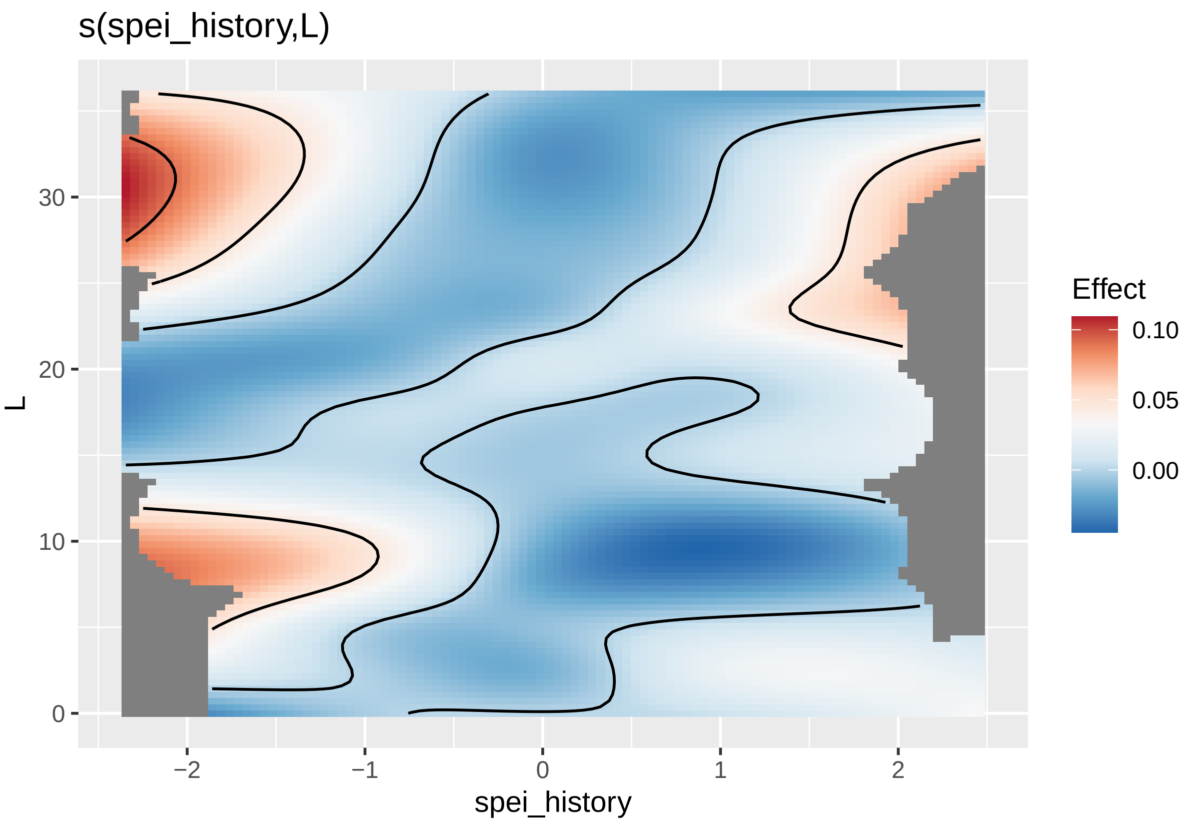

```
draw(g_cf_sub, select = "s(spei_history,L)", n_contour = 5)
```

```
## Warning: Removed 910 rows containing non-finite values (stat_contour).
```

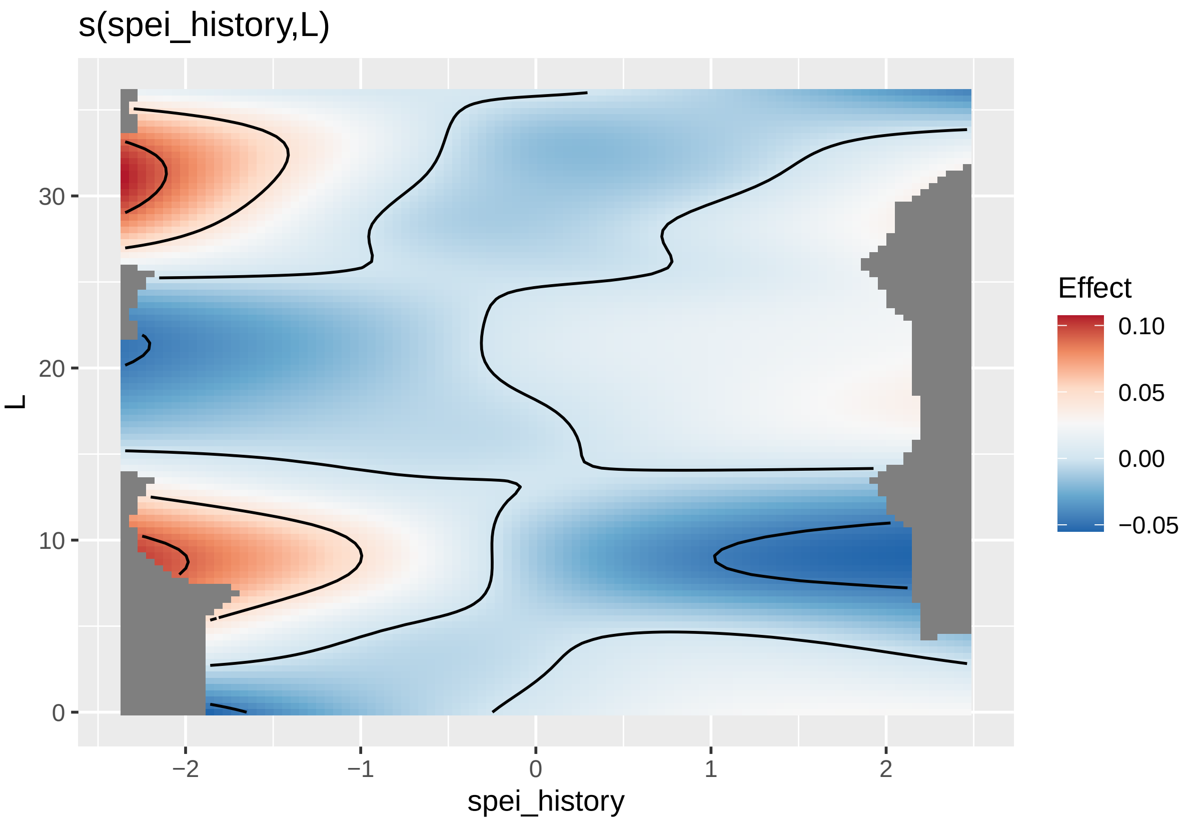

The shape of the smooth is also similar.

### Flowering

#### Fragments

##### Diagnostics

```
f_1ha_qres <- qresid(f_1ha)
par(mfrow = c(2,2))
#histogram
plot(density(f_1ha_qres))
#QQ plot
qqnorm(f_1ha_qres); qqline(f_1ha_qres)
#Fitted vs. residuals--binned residuals plot
binnedplot(fitted(f_1ha), residuals(f_1ha, type = "response"))
```

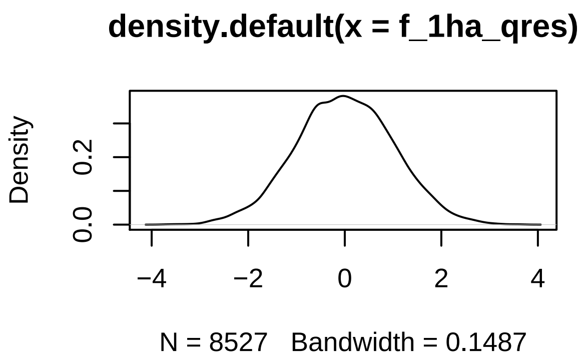

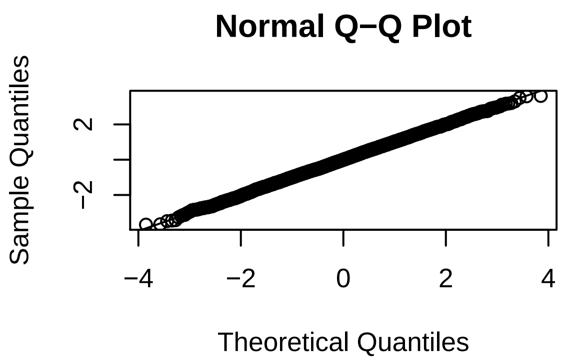

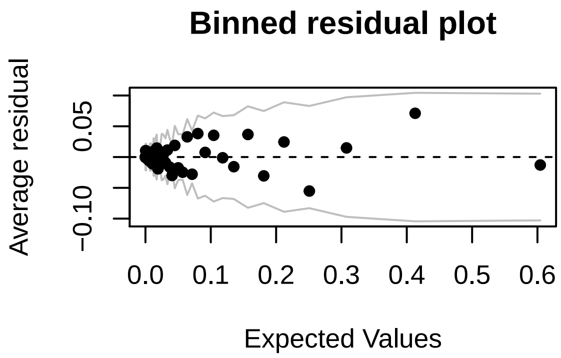

##### Basis dimensions

```
k.check(f_1ha)
```

```
##                      k'       edf   k-index p-value
## s(log_size_prev)     9  3.400302 0.9415327    0.04
## s(spei_history,L) 162 14.127281        NA      NA
## s(plot)              4  2.514545        NA      NA
```

Adequate k for `log_size_prev`

```
# looking for near zero edf
check_res_edf(f_1ha)
```

```
## # A tibble: 1 x 2
##   smooth                 edf
##   <chr>                <dbl>
## 1 te(spei_history,L)       0
```

Adequate k for crossbasis smooth.

#### Continuous Forest

##### Diagnostics

```r
f_cf_qres <- qresid(f_cf)
par(mfrow = c(2,2))
#histogram
plot(density(f_cf_qres))
#QQ plot
qqnorm(f_cf_qres); qqline(f_cf_qres)
#Fitted vs. residuals--binned residuals plot
binnedplot(fitted(f_cf), residuals(f_cf, type = "response"))
```

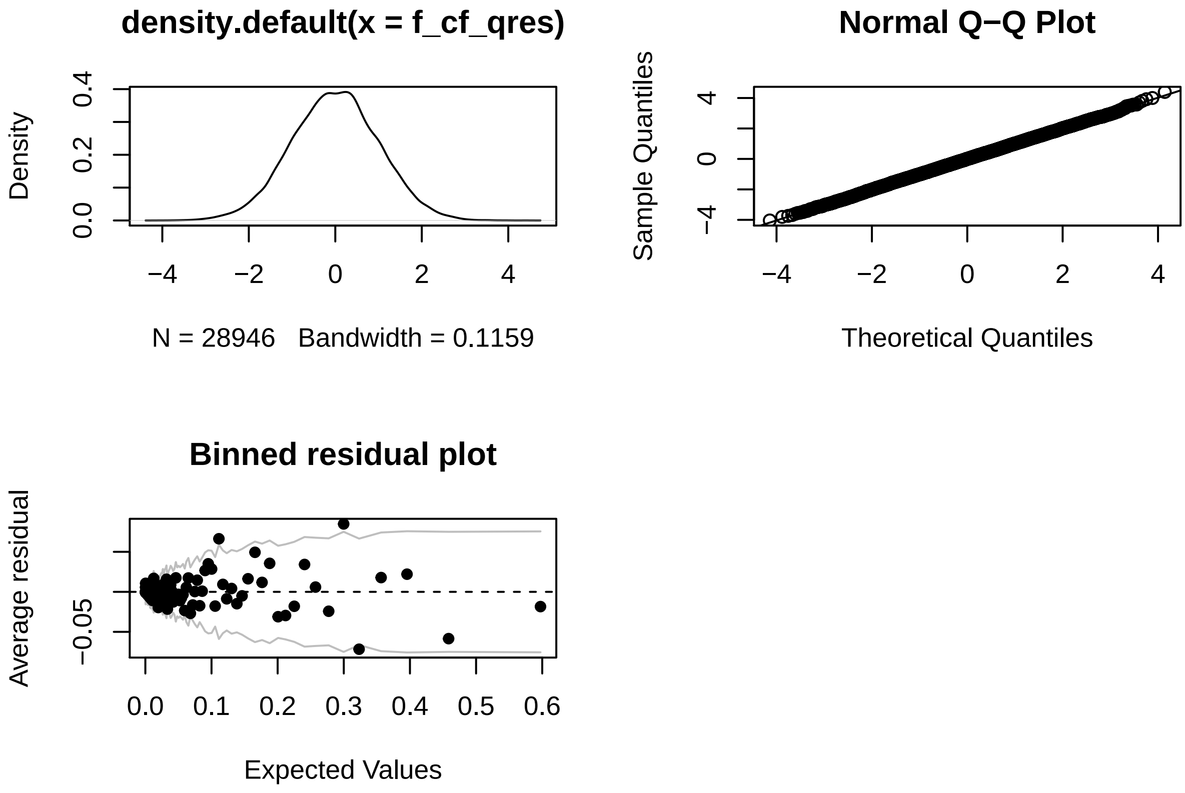

##### Basis dimensions

```r
k.check(f_cf)
```

```
##                     k'       edf   k-index p-value
## s(log_size_prev)     9  5.774695 0.9597626    0.19
## s(spei_history,L) 210 11.545081        NA      NA
## s(plot)              6  3.920380        NA      NA
```

Adequate k for `log_size_prev`

```
# looking for near zero edf
check_res_edf(f_cf)
```

```
## # A tibble: 1 x 2
##   smooth                edf
##   <chr>               <dbl>
## 1 te(spei_history,L)     1
```

Adequate k for crossbasis.

#### Effect of sample size

To check that differences are not purely due to sample size differences, particularly that lower edf in continuous forests is due to higher sample size.

```
summary(f_1ha)$edf[2]
```

```
## [1] 14.12728
```

```
summary(f_cf)$edf[2]
```

```
## [1] 11.54508
```

```
summary(f_cf_sub)$edf[2]
```

```
## [1] 11.3293
```

Similar edf for sub-sample

```
draw(f_cf, select = "s(spei_history,L)", n_contour = 5)
```

```
## Warning: Removed 968 rows containing non-finite values (stat_contour).
```

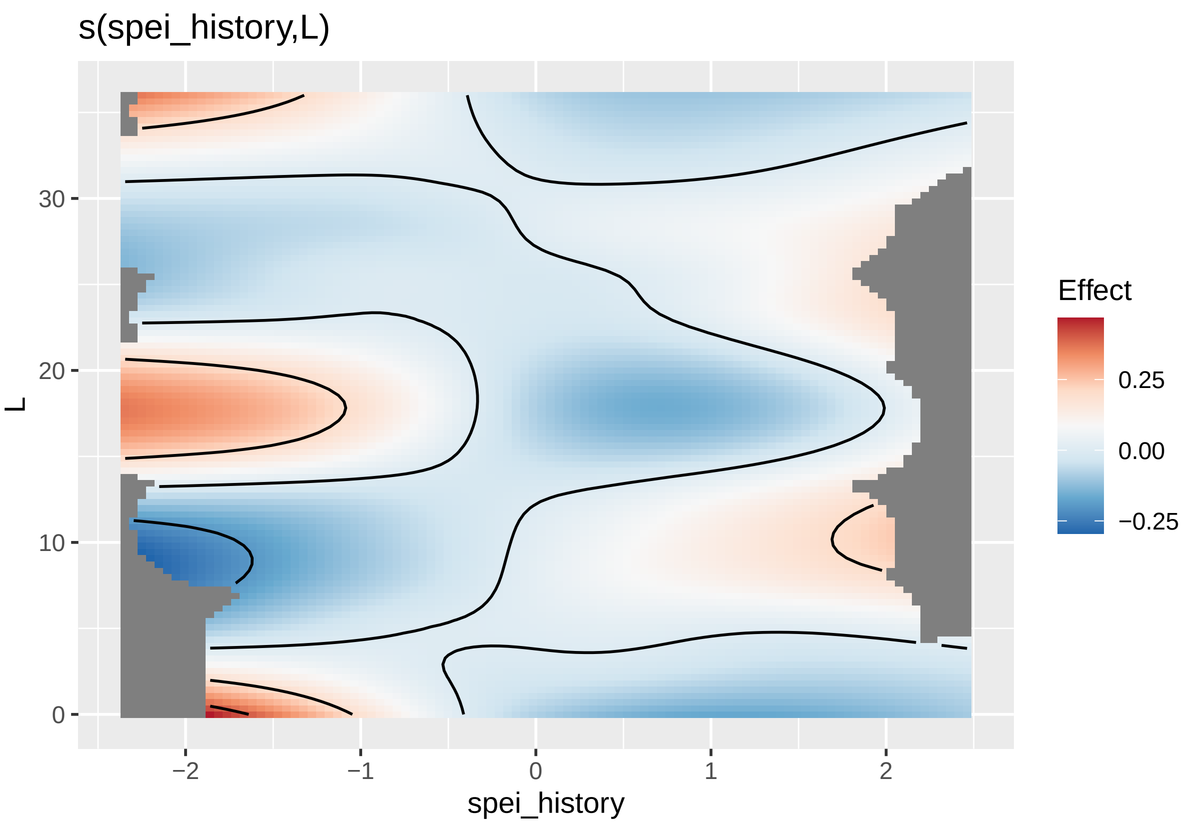

```r
draw(f_cf_sub, select = "s(spei_history,L)", n_contour = 5)
```

```
## Warning: Removed 910 rows containing non-finite values (stat_contour).
```

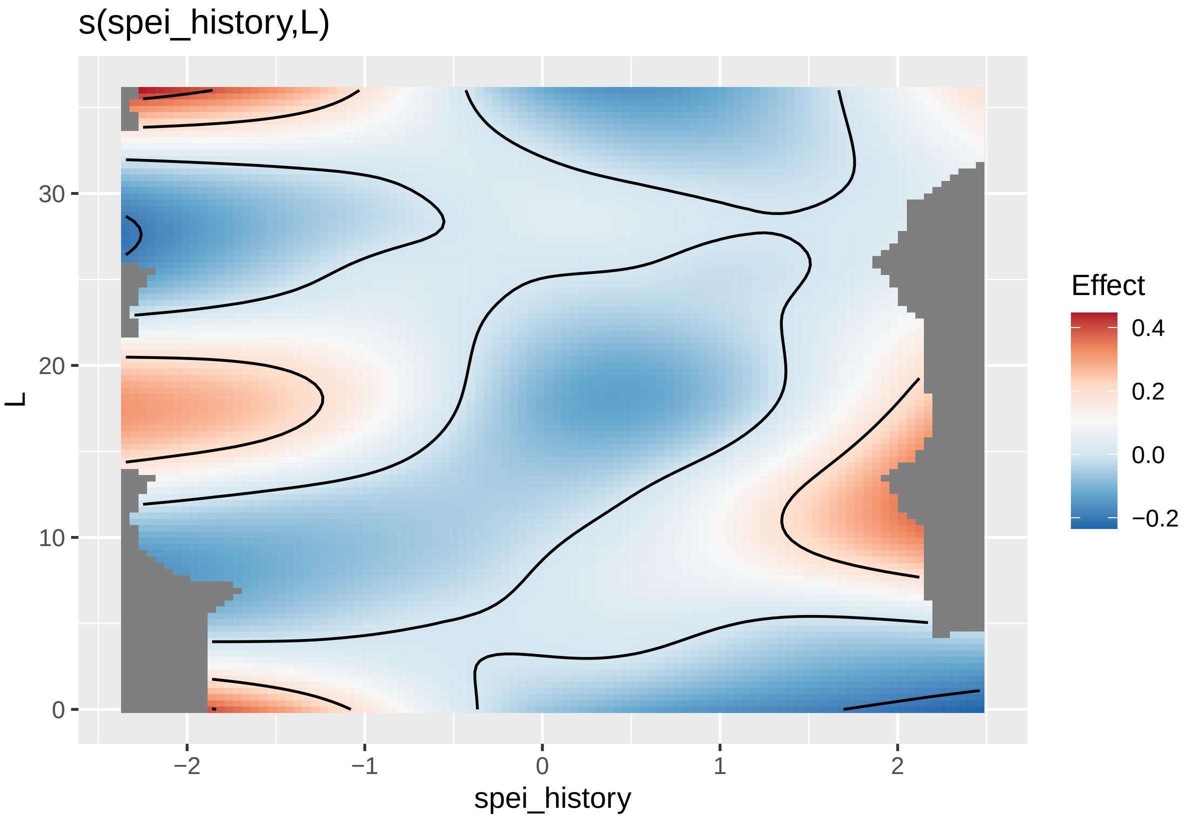

Similar, but essentially linear in SPEI dimension.

### Reproducibility

Reproducibility receipt

```
## datetime
Sys.time()
```

```
## [1] "2021-09-20 12:11:54 EDT"
```

```
## repository
if(requireNamespace('git2r', quietly = TRUE)) {
  git2r::repository()
} else {
  c(
    system2("git", args = c("log", "--name-status", "-1"), stdout = TRUE),
    system2("git", args = c("remote", "-v"), stdout = TRUE)
  )
}
```

```
## Local:    master /Users/scottericr/Documents/HeliconiaDemography
## Remote:   master @ origin (https://github.com/BrunaLab/HeliconiaDemography.git)
## Head:     [c920327] 2021-09-17: export fig for graphical abstract
```

```r
## session info
sessionInfo()
```

```
## R version 4.0.2 (2020-06-22)
## Platform: x86_64-apple-darwin17.0 (64-bit)
## Running under: macOS  10.16
##
## Matrix products: default
## BLAS:   /Library/Frameworks/R.framework/Versions/4.0/Resources/lib/libRblas.dylib
## LAPACK: /Library/Frameworks/R.framework/Versions/4.0/Resources/lib/libRlapack.dylib
##
## locale:
## [1] en_US.UTF-8/en_US.UTF-8/en_US.UTF-8/C/en_US.UTF-8/en_US.UTF-8
##
## attached base packages:
## [1] stats4    parallel  stats     graphics  grDevices datasets  utils
## [8] methods   base
##
## other attached packages:
##  [1] arm_1.11-2           lme4_1.1-27.1         Matrix_1.3-3
##  [4] MASS_7.3-54          performance_0.7.1     flextable_0.6.7
##  [7] pointblank_0.7.0.9000 conflicted_1.0.4     ggpointdensity_0.1.0
## [10] scales_1.1.1         pander_0.6.4          qqplotr_0.0.5
## [13] Hmisc_4.5-0          Formula_1.2-4         survival_3.2-11
## [16] lattice_0.20-44      readxl_1.3.1          colorspace_2.0-1
## [19] rmarkdown_2.7        statmod_1.4.36        latex2exp_0.5.0
## [22] gratia_0.6.0.9112    broom_0.7.6           patchwork_1.1.1
## [25] glue_1.4.2           bbmle_1.0.23.1        dlnm_2.4.5
## [28] mgcv_1.8-36          nlme_3.1-152          lubridate_1.7.10
## [31] janitor_2.1.0        tsModel_0.6           SPEI_1.7
## [34] lmomco_2.3.6         tsibble_1.0.1         forcats_0.5.1
## [37] stringr_1.4.0        dplyr_1.0.5           purrr_0.3.4
## [40] readr_1.4.0          tidyr_1.1.3           tibble_3.1.1
## [43] ggplot2_3.3.5        tidyverse_1.3.1       here_1.0.1
## [46] tarchetypes_0.2.0    targets_0.4.2         dotenv_1.0.3
##
## loaded via a namespace (and not attached):
##   [1] uuid_0.1-4          backports_1.2.1     systemfonts_1.0.1
##   [4] plyr_1.8.6          igraph_1.2.6        splines_4.0.2
##   [7] digest_0.6.27       htmltools_0.5.1.1   fansi_0.4.2
##  [10] magrittr_2.0.1      checkmate_2.0.0     memoise_2.0.0
##  [13] cluster_2.1.2       see_0.6.3           modelr_0.1.8
##  [16] officer_0.3.19      bdsmatrix_1.3-4     anytime_0.3.9
##  [19] jpeg_0.1-8.1        ggrepel_0.9.1       rvest_1.0.0
##  [22] haven_2.4.1         xfun_0.22           callr_3.7.0
##  [25] crayon_1.4.1        jsonlite_1.7.2      gtable_0.3.0
##  [28] DEoptimR_1.0-8      abind_1.4-5         mvtnorm_1.1-1
##  [31] DBI_1.1.1           Rcpp_1.0.6          isoband_0.2.4
##  [34] viridisLite_0.4.0   htmlTable_2.1.0     foreign_0.8-81
##  [37] htmlwidgets_1.5.3   httr_1.4.2          RColorBrewer_1.1-2
##  [40] ellipsis_0.3.2      farver_2.1.0        pkgconfig_2.0.3
##  [43] nnet_7.3-16         sass_0.3.1          dbplyr_2.1.1
##  [46] utf8_1.2.1          effectsize_0.4.4-1  labeling_0.4.2
```

```
##  [49] tidyselect_1.1.1    rlang_0.4.11        munsell_0.5.0
##  [52] blastula_0.3.2      cellranger_1.1.0    tools_4.0.2
##  [55] cachem_1.0.4        cli_2.5.0           generics_0.1.0
##  [58] ggridges_0.5.3      evaluate_0.14       fastmap_1.1.0
##  [61] yaml_2.2.1          goftest_1.2-2       processx_3.5.2
##  [64] knitr_1.33          fs_1.5.0            zip_2.1.1
##  [67] robustbase_0.93-7   mvnfast_0.2.5.1     xml2_1.3.2
##  [70] compiler_4.0.2      rstudioapi_0.13     png_0.1-7
##  [73] gt_0.2.2            reprex_2.0.0        clustermq_0.8.95.1
##  [76] bslib_0.2.4         stringi_1.6.2       parameters_0.13.0
##  [79] highr_0.9           ps_1.6.0            gdtools_0.2.3
##  [82] nloptr_1.2.2.2      commonmark_1.7      vctrs_0.3.8
##  [85] pillar_1.6.0        lifecycle_1.0.0     jquerylib_0.1.4
##  [88] insight_0.13.2      data.table_1.14.0   R6_2.5.0
##  [91] latticeExtra_0.6-29 bookdown_0.22       renv_0.13.2
##  [94] gridExtra_2.3       Lmoments_1.3-1      codetools_0.2-18
##  [97] boot_1.3-28         assertthat_0.2.1    rprojroot_2.0.2
## [100] withr_2.4.2         bayestestR_0.9.0    hms_1.1.0
## [103] grid_4.0.2          rpart_4.1-15        coda_0.19-4
## [106] minqa_1.2.4         snakecase_0.11.0    git2r_0.28.0
## [109] numDeriv_2016.8-1.1 base64enc_0.1-3
```
