## Appendix B for "Delayed effects of climate on vital rates lead to demographic divergence in Amazonian forest fragments"

Eric R. Scott

María Uriarte

Emilio M. Bruna

20 September, 2021

### Climate Normals for BDFFP

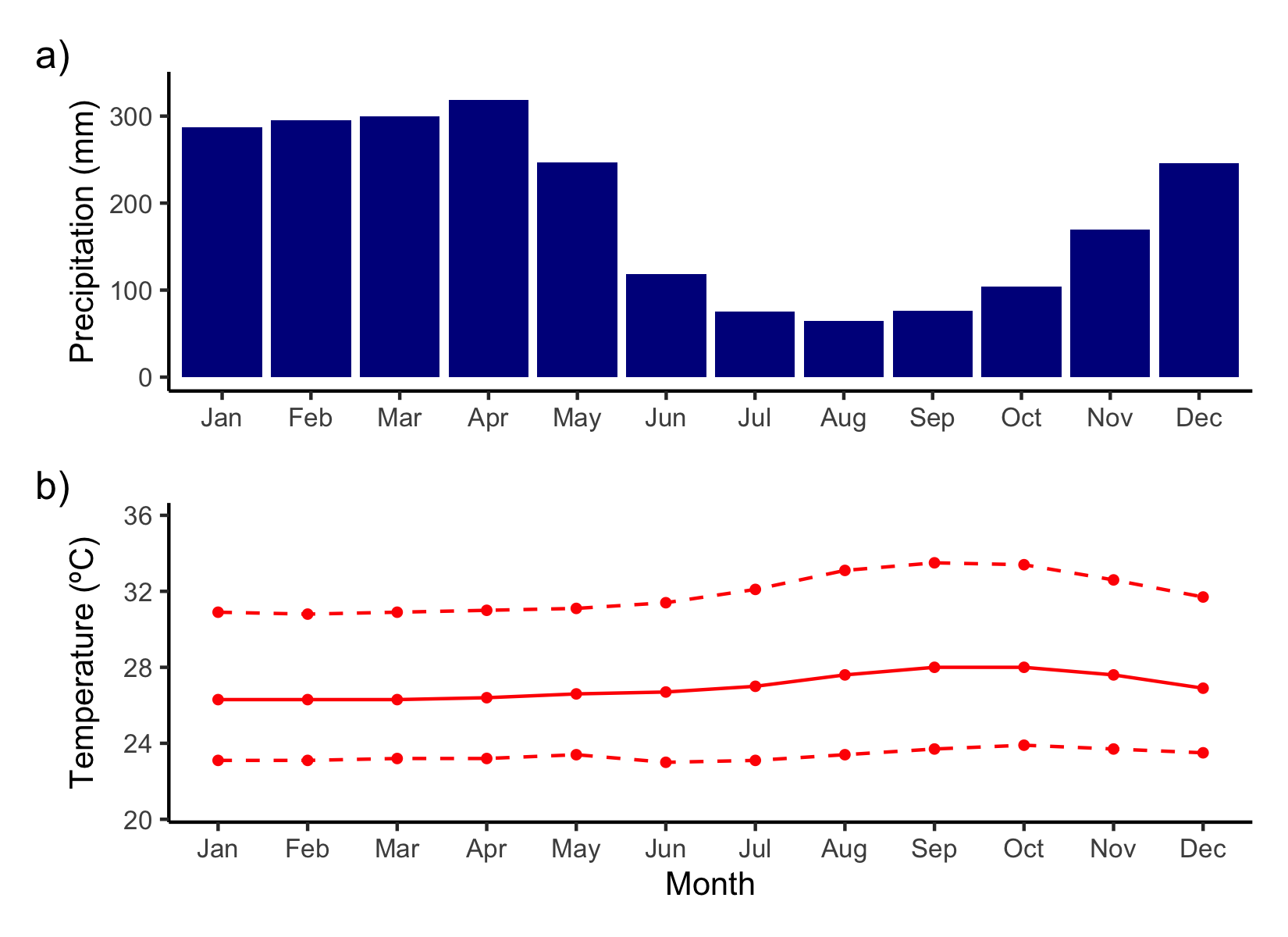

Figure S1. Weather normals for Manaus, Brazil (3º6’S, 60º1’W) for 1981–2010. Precipitation (a) shows a marked dry seasons from June through October. Temperature (b) varies less throughout the year. Mean monthly temperature is shown in the solid red line and monthly minimum and maximum temperatures are shown with the lower and upper dashed lines, respectively. Data from Brazilian National Institute of Meterology (INMET).

### Comparison of SPEI calculated from different data sources

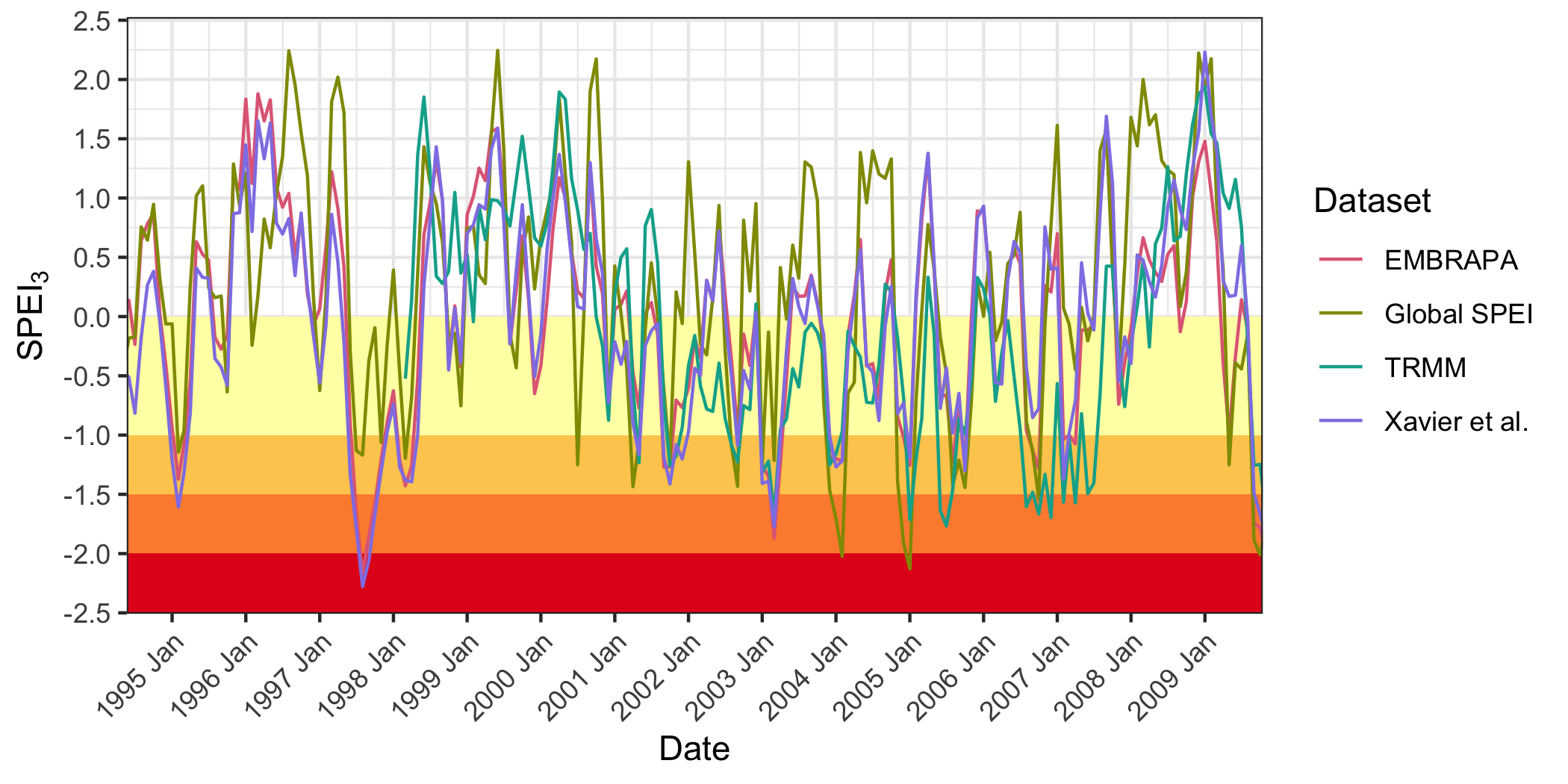

Figure S2. Three-month SPEI at BDFFP calculated using 4 datasets. “EMBRAPA” (red) is the average of SPEI calculated using precipitation and evapotranspiration from two weather stations near BDFFP—Estacao Rio Preto da Eva ( -2.65º, -59.65º) and Estacao Manaus (-2.65º, -60.35º). The data was accessed from the EMBRAPA Infoclima data portal (<https://www.cnpaf.embrapa.br/infoclima/>). “Global SPEI” (green) is from the SPEI global database(Beguería et al., 2010; Vicente-Serrano et al., 2010). “TRMM” (blue) SPEI uses precipitation data from TRMM ((TRMM), 2011) and evapotranspiration data from Xavier et al. (2016) to calculate 3-month SPEI. Note that TRMM data only goes back to 1998. Xavier et al. (2016) is the SPEI source described in the main text. Drought categories mild, moderate, severe, and extreme drought are represented by yellow, goldenrod, orange, and red panels, respectively.

Table S1. Comparison of SPEI values and drought categories across datasets for major drought events in the Amazon. Mild, moderate, severe, and extreme drought are represented by yellow, goldenrod, orange, and red, respectively. Cause indicates whether the drought was caused by the El Niño Southern Oscillation (ENSO) or Atlantic Sea Surface Temperatures (AMO). The lowest SPEI value for the year identified by each dataset is reported, along with the month that minimum value occurred in. Datasets are described in the legend for Figure S2.

|  |  | Lowest SPEI for the year (month) | | | |
| --- | --- | --- | --- | --- | --- |
| year | cause | Global SPEI | Xavier et al. | EMBRAPA | TRMM |
| 1997 | ENSO^a^ | -1.17 (Aug) | -2.28 (Aug) | -2.17 (Aug) | – |
| 2002 | ENSO | -1.43 (Sep) | -1.09 (Sep) | -0.96 (Sep) | -1.23 (Sep) |
| 2004 | ENSO | -2.02 (Feb) | -1.27 (Jan) | -1.22 (Feb) | -1.16 (Jan) |
| 2005 | AMO^b^ | -2.13 (Jan) | -1.3 (Oct) | -1.25 (Jan) | -1.77 (Jul) |
| 2006 | ENSO | -1.53 (Oct) | -0.85 (Sep) | -1.28 (Oct) | -1.7 (Dec) |
| 2009 | ENSO | -2.01 (Oct) | -1.88 (Nov) | -2.04 (Nov) | -1.72 (Nov) |
| 2010 | AMO^c^ | -0.76 (May) | -1.12 (Mar) | -1.4 (Mar) | -1.1 (May) |
| ^a^McPhaden 1999; ^b^Marengo et al. 2008; Zeng et al. 2008; ^c^Lewis et al. 2011 | | | | | |

### Choice of variable for plant size

Table S2. Using data from transplant experiments of *Heliconia acuminata* (Bruna, 2021; Bruna et al., 2002) we fit linear regressions with log(leaf area) as the response variable and either log(height), log(shoot number) or log(shoot number x height) as the predictor. Using the log of size (shoot number $\times$ height) as a predictor gives the lowest AIC, BIC, root mean squared error (RMSE) and highest $R^{2}$ indicating that it is the best proxy for predicting log(leaf area) as a response.

| Model | dAIC | dBIC | R2 | RMSE |
| --- | --- | --- | --- | --- |
| log(leaf area) ~ log(shoot number x height) | 0.00 | 0.00 | 0.67 | 0.342 |
| log(leaf area) ~ log(height) | 72.77 | 72.77 | 0.63 | 0.366 |
| log(leaf area) ~ log(shoot number) | 459.13 | 459.13 | 0.24 | 0.521 |

### Size plots with raw data

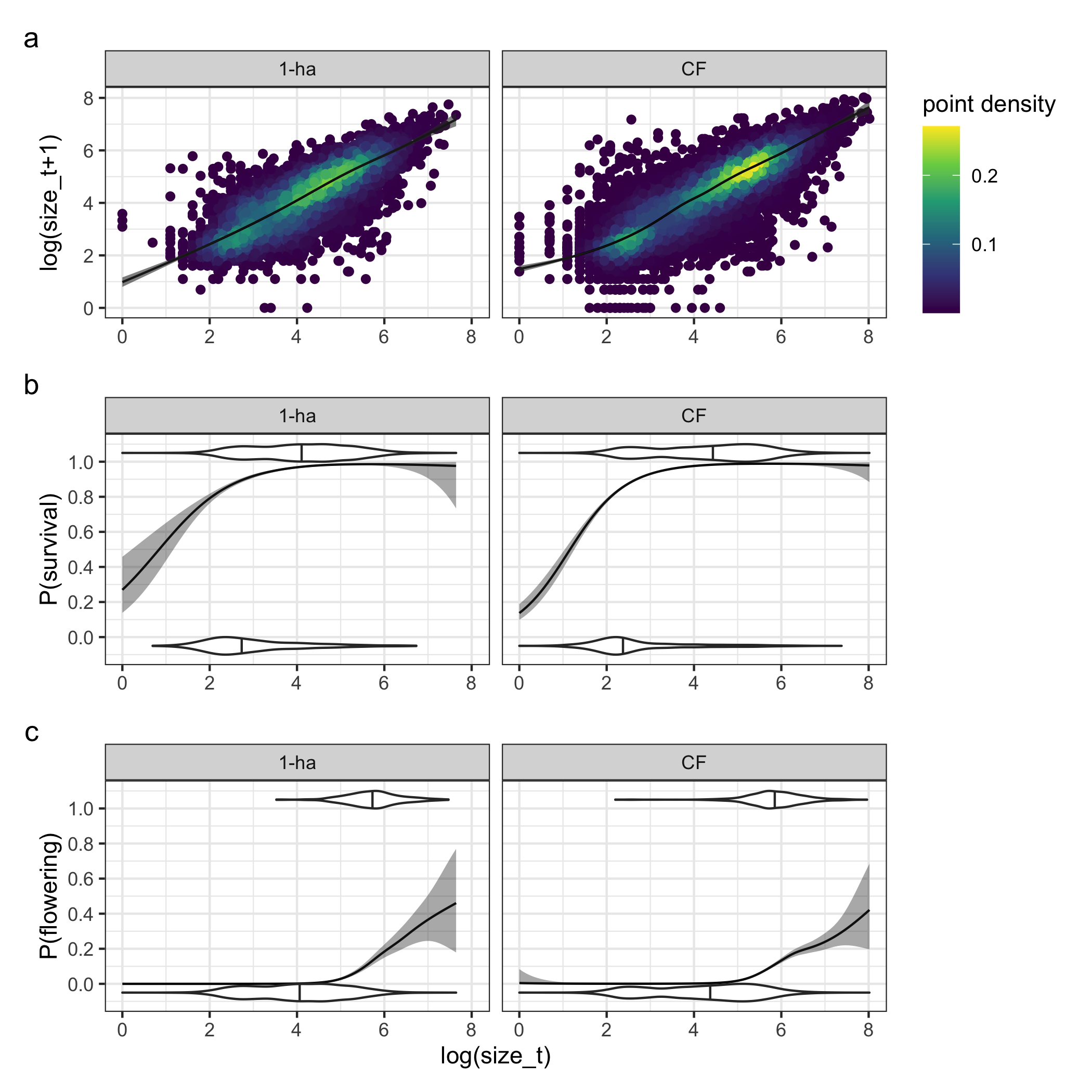

Figure S3. The curves and confidence intervals plotted here are the same as those in Figure 3a–c but separated by habitat (1-ha fragments in left panel, continuous forest plots in right panel) and with raw data superimposed. For size (a), raw data are plotted as points with color representing point density to assist in visualizing overplotted data. For survival (b) and flowering (c), raw data (survived or flowered = 1, died or didn’t flower = 0) are shown using violin plots with the median marked by a vertical line.
